# An organoid and multi-organ developmental cell atlas reveals multilineage fate specification in the human intestine

**DOI:** 10.1101/2020.07.24.219147

**Authors:** Qianhui Yu, Umut Kilik, Emily M. Holloway, Yu-Hwai Tsai, Angeline Wu, Joshua H. Wu, Michael Czerwinski, Charlie Childs, Zhisong He, Ian A. Glass, Peter D. R. Higgins, Barbara Treutlein, Jason R. Spence, J. Gray Camp

## Abstract

Human intestinal organoids (HIOs) generated from pluripotent stem cells provide extraordinary opportunities to explore development and disease. Here, we generate a single-cell transcriptome reference atlas from HIOs and from multiple developing human organs to quantify the specificity of HIO cell fate acquisition, and to explore alternative fates. We identify epithelium-mesenchyme interactions, transcriptional regulators involved in cell fate specification, and stem cell maturation features in the primary tissue that are recapitulated in HIOs. We use an HIO time course to reconstruct the molecular dynamics of intestinal stem cell emergence, as well as the specification of multiple mesenchyme subtypes. We find that the intestinal master regulator CDX2 correlates with distinct phases of epithelial and mesenchymal development, and CDX2 deletion perturbs the differentiation of both intestinal epithelium and mesenchyme. Collectively our data provides a comprehensive and quantitative assessment of HIO development, and illuminates the molecular machinery underlying endodermal and mesodermal cell fate specification.

## Introduction

Complex multicellular culture systems that accurately recapitulate human physiology can be used to understand human development and model human disease. Human pluripotent stem cell (hPSC)-derived intestinal organoids (HIOs) are an attractive model system to study the intestine because they can be genetically manipulated, observed in controlled in vitro environments, and derived from individuals with diverse genetic backgrounds [1]. HIOs also recapitulate the earliest phases of organ specification and complex tissue formation, enabling exploration of human development at time points that are generally considered inaccessible. Generating HIOs relies on directed differentiation, a process that uses temporal manipulation of key developmental signaling pathways via exogenously supplied growth factors and small molecules to mimic intestinal organogenesis in a step-by-step manner [2, 3]. For HIOs, definitive endoderm and mesoderm are co-differentiated from pluripotency and are further differentiated into a mid/hindgut fate, resulting in the aggregation of three dimensional (3D) spheroids that bud from an adherent cell monolayer. Spheroids are subsequently collected and embedded in a 3D matrix with pro-intestinal molecules [3–5] to promote intestinal growth, morphogenesis, and differentiation. The resulting HIOs consist of a polarized, columnar epithelium that is surrounded by mesenchymal cells. After transplantation into immunocompromised mice, the epithelium matures and gives rise to a crypt-villus morphology. Stem cells are located in proliferative domains at the crypt base with enterocytes, goblet, and enteroendocrine cells distributed along the crypt-villus axis [6–8]. The HIO culture system has been used over the past years as a model to study human intestinal development [1, 6–14] and disease [1, 15–22]. Nevertheless, cellular diversity within HIOs, especially that of HIO mesenchyme, has not yet been comprehensively resolved.

Quantitative comparisons to primary human tissues are needed to fully understand the potential and limitations of HIOs. There have been single-cell transcriptome surveys that illuminate cell composition and epithelial differentiation programs in mouse [23–26] and human [27] intestinal epithelial organoids derived from adult stem cells. Reference atlases are also being established from developing and adult primary human intestinal tissues [28–31] to identify molecular features underlying intestine development. Single-cell maps of gastrulation and other early stages of organ specification and development in mice can provide insight into processes that are difficult to explore in human primary tissues [32, 33]. Yet, how well HIOs recapitulate the in vivo cell type features remains to be determined. Furthermore, it was shown that hPSC-derived cells and tissues have prolonged inherent plasticity after differentiation in vitro, resulting in off-target cell types in culture [34]. This observation raises the necessity of comparing HIOs to primary tissues of intestine and other organs to assess whether or not HIOs give rise to non-intestinal cell types.

Here we used single-cell messenger RNA sequencing (scRNA-seq) to decipher cellular composition in HIOs during in vitro differentiation and growth and after in vivo maturation in an immunocompromised mouse. We further generated a multi-organ developmental atlas covering endoderm-derived organs along the human gastrointestinal (GI) tract, including esophagus, stomach, lung, liver, and intestine (duodenum, jejunum, ileum and colon), from multiple time points and biological specimens. We used this atlas to benchmark cellular composition and molecular profiles within in vitro and in vivo HIOs. By comparing HIOs to the developing human reference, we quantified the similarity between primary tissue counterparts and identified off-target cell types. We use these resources to delineate the transcription factors and molecular mechanisms that are specific to the mesenchyme of each organ and use receptor-ligand pairing analysis to gain insights into signaling between mesenchymal subpopulations and the adjacent epithelium. We use HIOs to track the early steps of intestinal epithelial stem cell development, as well as the emergence of mesenchymal subpopulations. We find that intestinal master regulator CDX2 is required not only for intestinal epithelial cell fate specification, but also necessary for proper specification of the gut-associated mesenchyme. These data provide a quantitative assessment of HIO development and illuminate the molecular machinery underlying endodermal and mesodermal cell fate specification.

## Results

### Deconstructing cell type heterogeneity in human intestinal organoids (HIOs)

We used single-cell transcriptomics (scRNA-seq) to analyze cellular heterogeneity in HIOs grown in vitro for 4-weeks, as well as HIOs 4- and 8-weeks after in vivo transplantation (tHIOs) into the kidney capsule of an immunocompromised mouse host (Figure 1A; Table S1). Consistent with previous work [1, 7], immunohistochemistry of both HIOs and tHIOs revealed that VIM+ mesenchyme surrounds an ECAD+ epithelium and the relatively immature in vitro grown HIO morphologically matures into a crypt-villus architecture containing differentiated epithelial cell types upon transplantation (Figure 1B). ScRNA-seq data revealed multiple molecularly distinct clusters (c) of mesenchymal (c1-4) and epithelial (c5-8) cells in the 4-week HIO in vitro (Figures 1C, 1D, S1A-S1D; Table S2). Two epithelial clusters (c5, MKI67+; c6, LGALS4+/CDH17+) express the canonical intestinal master regulator, CDX2, suggesting midgut/hindgut specification, which accounts for 69.2% (2,186/3,159) of sequenced epithelial cells in the 4-week in vitro HIO dataset. However, two clusters, accounting for 30.8% (973/3,159) of analyzed epithelial cells, do not appear to be committed to the intestinal lineage as one cluster does not express CDX2 (c8, SOX2+/MIR205HG+), and the other (c7, CLDN18+/ANXA10+) co-expresses low levels of CDX2 along with the foregut markers SOX2 and MUC5AC (Figure 1D). Epithelial precursor cells (c6) co-express intestinal epithelial membrane trafficking genes CDH17 [35] and LGALS4 [36], however, the majority of these cells are FABP2-/SI- (sucrase-isomaltase) (96%) or MUC2-/SPINK4- (91%), suggesting absorptive enterocytes and goblet cells have not yet differentiated at this time point. We note that out of 3,159 sequenced epithelial cells, the stem cell marker LGR5 was detected in only 56 cells (1.8%), and these LGR5+ cells do not form a distinct cluster within the UMAP cell embedding, suggesting the absence of LGR5+ stem cells at this stage. This result is consistent with recent findings on the early-stage mouse and human intestine revealing that LGR5+ stem cells were absent before intestine villus formation [31, 37], and started to emerge at around 10 post-conception weeks (PCW) in humans.

**Figure 1.**
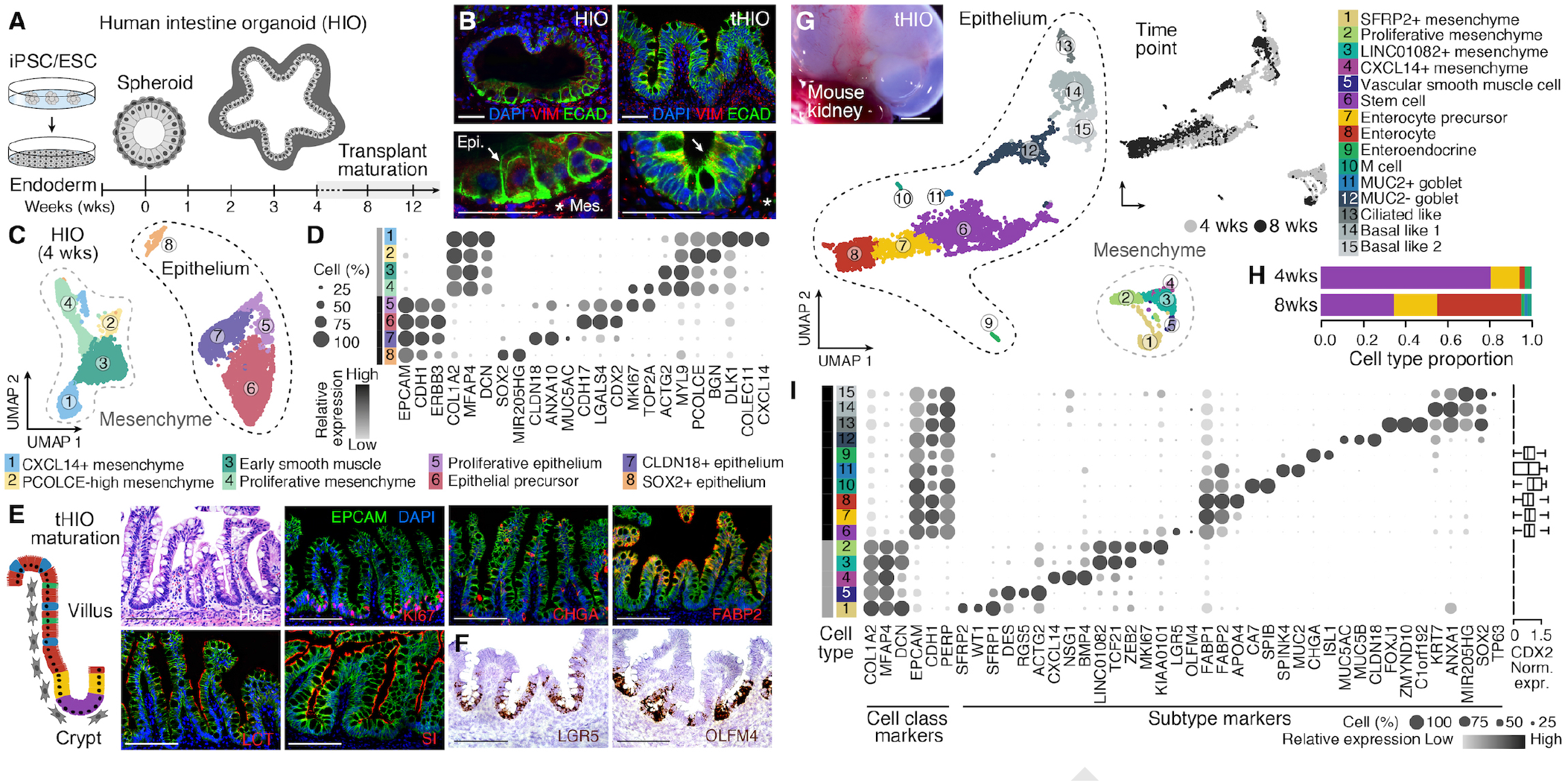
Deconstructing cell type heterogeneity in human intestinal organoids (HIOs). A) Schematic of HIO development. B) Immunofluorescence (IF) stainings showing expression of mesenchyme (VIM, red, star) and epithelium (ECAD, green, arrow) markers in day 30 HIO and 10-week tHIO. Nuclei stained with DAPI (blue). Bottom is a magnification of the top image. Scale bars, 200 μm. C) UMAP embedding of single-cell transcriptomes from 4-week HIO colored and numbered by cluster. D) Dot plot showing HIO marker gene expression levels. Dot color and size are proportional to averaged cluster expression and proportion of cells expressing, respectively. E) HIO histological analysis 10 weeks after transplantation into immunocompromised mouse kidney (tHIO). Left, schematic of epithelial cell types. Hematoxylin and Eosin, HE; Lactase, LCT; Marker Of proliferation Ki-67, Ki67; Sucrase-isomaltase, SI; Chromogranin A, CHGA; Fatty acid-binding protein 2, FABP2. Scale bar, 200 μm. F) RNA in situ hybridization of stem cell marker genes in 10-weekt HIOs. Leucine-rich repeat-containing G-protein coupled receptor 5, LGR5; Olfactomedin 4, OLFM4. Scale bars, 200 μm. G) UMAP embedding of single-cell transcriptomes from tHIO (4 and 8 weeks after transplantation) colored and numbered by cluster. Top left, 10-week tHIO in the mouse kidney capsule. Scale bar represents 1 mm. Top right, UMAP embedding with cells colored by time point. H) Stacked bar plot showing epithelial cell type composition in 4- and 8-week tHIO for intestinal cells (clusters 6-11). Color scheme as in panel G. (I) Dot plot of cluster marker gene expression. Dot color and size are proportional to average cluster expression and proportion of cells expressing, respectively.

Transplanted HIOs exhibited hallmarks of maturation similar to what is observed in the developing human intestine [6–8]. Immunohistochemistry confirms a polarized epithelium with proliferative MKI67+ stem cells located in a crypt base, and differentiated secretory lineages (CHGA+ enteroendocrine, MUC2+ goblet cells) and absorptive enterocytes showing apical expression of the brush border enzymes sucrase isomaltase (SI) and lactase (LCT), and apical sodium–bile acid transporter (ASBT, SLC10A2), emerging along the villus epithelium (Figures 1E and S2A). Based on the molecular profiles from the single-cell transcriptome data (Figure 1G; Table S2), we observed intestinal stem cells (ISCs) (c6, LGR5+/OLFM4+), enterocyte precursors (c7, FABP2+/APOA4 low), enterocytes (c8, FABP2+/APOA4+), enteroendocrine cells (c9, CHGA+/ISL1+), M cells (c10, SPIB+/CA7+), and goblet cells (c11, MUC2+/SPINK4+) (Figures 1F, 1G and S2A-S2F; 66.0%, 2,100/3,182 of tHIO epithelial cells). We also detected additional epithelial clusters (c12-15) that lacked the expression of canonical small intestinal marker genes (e.g. CDX2) (Figures 1H, 1I, S2G and S2H), and shared genes expressed in lung multiciliated and basal cells. These off-target clusters account for 34.0% (1,082/3,182) of sequenced epithelial cells. In addition, we find multiple mesenchymal clusters (c1-5), which are analyzed in more depth in later figures. We note that in contrast to the 4-week in vitro HIO, the LGR5+ ISC pool has expanded to 21.4% (680/3,182) of the epithelial cells sequenced in this tHIO dataset (Figure S2C). Together, this data suggests that the majority of cells within the HIO are specified to an intestinal fate in vitro and in vivo, and differentiated intestinal epithelial cells emerge after transplantation.

### Mapping to a multi-organ reference atlas reveals the specificity of tHIO cell fates

We next sought to quantify the similarity of tHIO epithelial and mesenchymal cells to primary human tissue, and determine the identity of off-target cells. We established a developing human multi-organ cell atlas by integrating single-cell transcriptomes from lung, liver, esophagus, stomach, small intestine (duodenum, jejunum, ileum), and colon (155,232 cells total) with an age distribution spanning 7 to 21 PCW (Figure 2A; Tables S1 and S2; Key Resources Table) [14, 30, 38]. We note that for some of the time points, we have sequenced multiple organs from the same human specimens (Figure S3C). We integrated all of the developing human scRNA-seq data using Cluster Similarity Spectrum (CSS) [39] to remove confounding effects of random or technical differences between samples. High-level clustering resolved multiple major cell classes, including epithelial, mesenchymal, immune, endothelial, neuronal, and erythroid populations (Figures 2B, 2C and S3A-S3D). Each class can be further subdivided into molecularly distinct clusters, which vary in their organ proportion across the clusters, and we note that epithelial and mesenchymal populations are diverse and have organ-specific features (Figure S3D). This multi-organ single-cell transcriptome reference atlas can be used to identify inter-lineage signaling and transcriptional programs that are specific to cell populations within each organ (Figures S3E and S3F).

**Figure 2.**
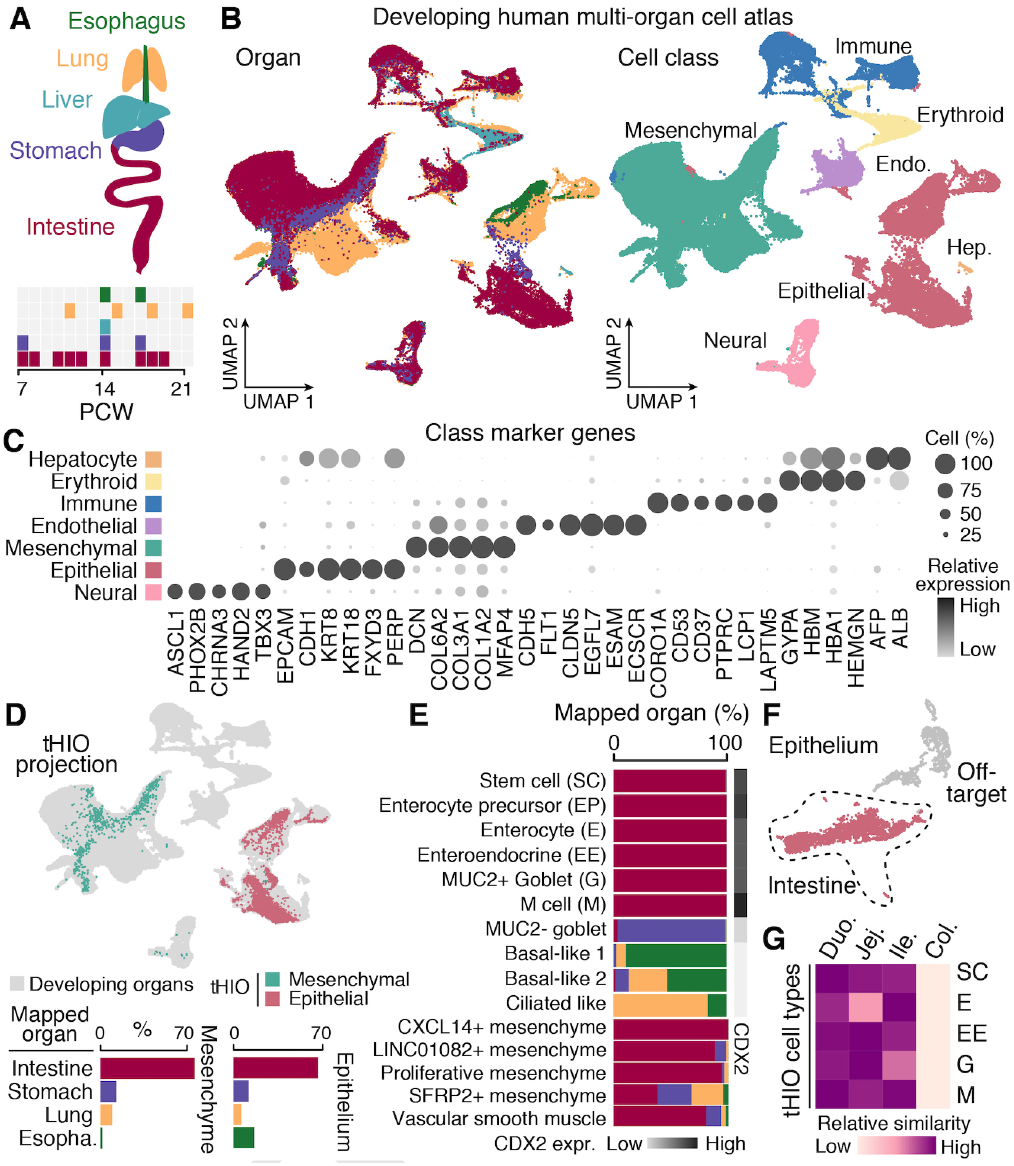
Mapping to a human multi-organ reference atlas reveals the specificity of tHIO cell fates. A) Schematic shows the organs and developmental time points (post-conception weeks, PCW) profiled by scRNA-seq in this study (155,232 cells total). B) UMAP embedding of scRNA-seq data colored by organ (left) and cell class (right). C) Dot plot shows class-level marker gene expression. Dot color and size is proportional to average cluster expression and proportion of cells expressing, respectively. D) Top, 8-week tHIO cells are projected onto the reference atlas embedding using UMAP transformation. Bottom, proportion of tHIO cells mapping to mesenchyme (left, teal) or epithelium (right, rose) of each organ. E) Stacked bar plot shows the proportion of cells in each tHIO cell type mapping to reference atlas organs. Sidebar shows CDX2 expression. F) tHIO UMAP embedding highlighting intestinal (rose) and off-target epithelial cells (grey). G) Heatmap showing relative Spearman’s correlation coefficients between tHIO and different intestine regions for each epithelial subtype. Only developing specimens with more than three intestine regions (N = 3) were used for comparison. Duodenum, Duo; Jejunum, Jej; Ileum, Ile; Colon, Col.

Our major goal was to use the atlas as a high-dimensional search space to determine the potential correspondence between tHIO cells and their in vivo counterparts. We projected the tHIO cells to the reference two-dimensional (2D) latent space and used Euclidean distance to search for the most similar cell type. We found that approximately 70% of tHIO epithelial and mesenchymal cells map to the developing human intestine (Figure 2D). For each tHIO cluster, we determined the proportion of cells mapped to each developing organ cell type cluster, and found that tHIO epithelial subtypes that are mapped to the intestine have high CDX2 expression compared to the cells mapping to non-intestine clusters (Figures 2E and 2F). To interrogate intestine region identity of tHIOs, we extracted epithelial cells of the developing intestine regions, including duodenum, jejunum, ileum and colon, and performed sub-clustering and cell type annotation (Figures S4A-S4D). We compared the tHIO intestine epithelial subtypes to their counterparts in various developing intestine regions, and found that tHIO cells are more similar to the small intestine than colon (Figures 2F and 2G; Table S2). Regional identity within the small intestine was not clearly resolved in primary tissue and tHIO from our data (Figures 2G and S4E). Taken together, tHIOs contain diverse epithelial and mesenchymal cells, most of which map to the developing small intestine relative to other organs.

### Integrating HIO and primary duodenum data enables tracking of molecular transitions during human small intestinal stem cell maturation

We next generated a single-cell transcriptome reference of adult duodenum (Fig. S5A-S5C; Table S2), and compared tHIO, developing, and adult epithelial subtypes to quantify tHIO epithelial cell maturation, and identify molecular features specific to different stages of small intestinal maturation. In an integrated UMAP embedding, we found that developing duodenum and tHIO stem cell-to-enterocyte differentiation trajectories were distinct from the adult (Figure 3A). Indeed, we found that each tHIO cell type was more similar to the developing primary counterpart compared to the adult (Figures 3B, S5D and S5E). Notably, we found the tHIO and developing duodenum stem cells to be highly similar, and molecularly distinct from the adult state (Figure 3C; Supplemental Videos). We performed differential gene expression analysis between the developing and adult ISC populations, and identified genes that were significantly enriched in developing or adult stem cells (Figure 3D; Table S3). Gene Ontology (GO) enrichment analysis of genes that were highly expressed in developing ISCs revealed slight biological process enrichments in nitric oxide regulation (argininosuccinate synthase 1, ASS1; clusterin, CLU; endothelin 1, EDN1), TGF-β signaling (FOS, JUN, ID1) and transcriptional regulation (ARID3A, FOS, AP-1, JUN, JUNB, EGR1) (Table S3). We also noted enriched expression of the transcription initiation factor EIF3E in the developing ISCs (Figure 3D), and EIF3E has been linked to cancer causing chromosomal rearrangements involving RSPO2 [40, 41]. We confirmed that the ISC marker, olfactomedin 4 (OLFM4) [42], is detected in the tHIO and and developing ISCs, but is expressed at significantly higher levels in the adult ISCs. Other examples of adult ISC-enriched genes include islet of Langerhans regenerating protein 1A (REG1A) and polymeric immunoglobulin receptor (PIGR).

**Figure 3.**
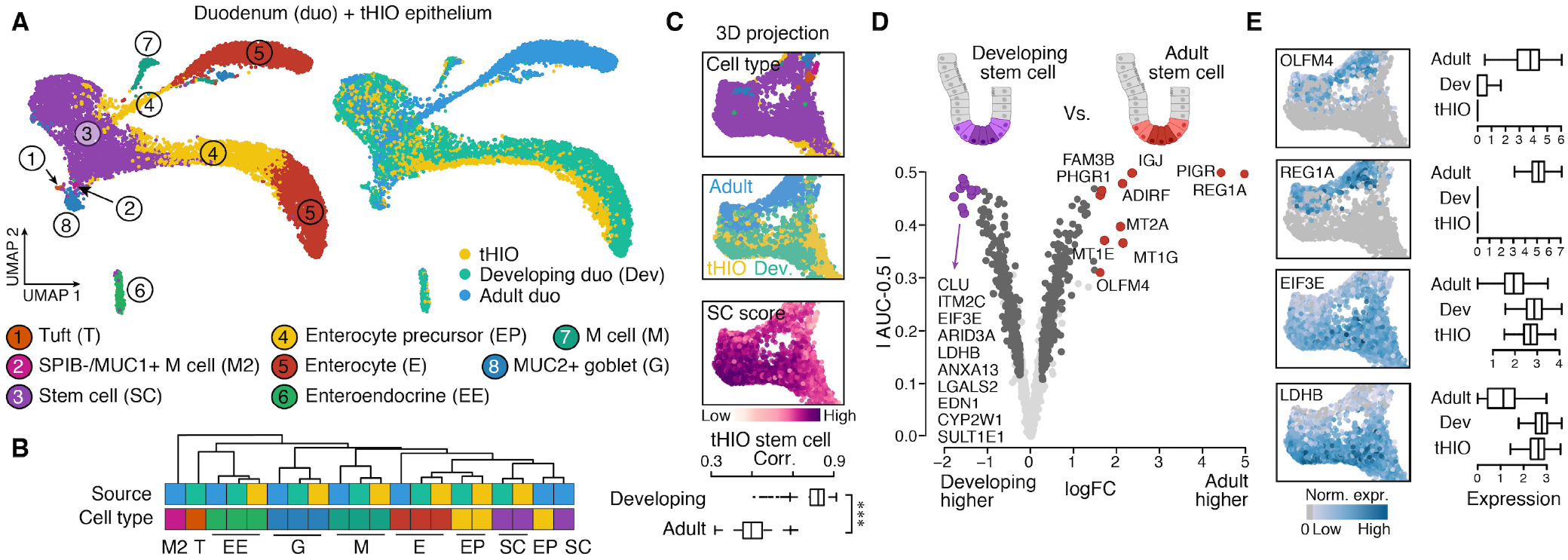
Developing human duodenum and tHIO epithelial stem cells are distinct from the adult state. A) Integrated UMAP embedding of tHIO, developing, and adult duodenum epithelial scRNA-seq data sets, with plots colored by cell type (left) or source (right). B) Hierarchical clustering of average transcriptome correlations between tHIO, developing, and adult duodenum cell types, with top sidebar coloring by tissue source. C) Inset of 3D UMAP embedding highlighting the stem cells colored by cell type (top), source (middle), and stem cell score (bottom). See also Supplemental Videos for complete view of the 3D UMAP embedding. Stem cell scores were calculated as cumulative z-transformed expressions of the union set of stem cell markers identified in the developing or adult tissue. Boxplot shows the distribution of Spearman’s correlation coefficients of tHIO stem cells compared to the developing or adult stem cells (***, Wilcoxon rank-sum test, nominal P < 0.0001). D) Volcano plot shows genes with higher expression in the developing (left, purple) or adult (right, red) ISCs. X-axis presents log-transformed expression fold change in adult versus developing duodenum; y-axis presents absolute difference between area under receiver operator (auROC, AUC) and 0.5, which was used to quantify the effectiveness of using a gene to classify two groups. E) Feature plots (left) and boxplot (right) show expression distributions of differentially expressed genes in adult, developing, and tHIO stem cells.

Notably, for genes that were differentially expressed between developing and adult ISCs, tHIOs consistently showed the developing intestine expression pattern (Figure 3E). This analysis shows that tHIO stem cells are more similar to the developing intestine than adult counterparts, and also reveals features that distinguish developing and adult intestine cell states.

Given the strong similarity of developing intestine and tHIO cell states, we utilized HIOs to explore early stages of human intestine development at time points that are otherwise inaccessible. We analyzed single-cell transcriptomes across a time course from endoderm induction through 30 days of in vitro HIO differentiation to illuminate the molecular transitions that lead to intestinal epithelial stem cell specification (Figure 4A). We constructed a force-directed k-nearest neighbour (kNN) graph [43] to visualize the temporal progression of cell fate acquisition (Figures 4B and S6A; Table S2). Given that we do not have human reference time points for the early differentiation events in HIOs, we compared the in vitro organoid time course to a mouse gastrulation reference atlas [33] to help annotate each cluster (Figures S6B-S6D). This analysis revealed that cells in the HIO early time point clusters (‘day 0’- spheroid, c4/c13) expressed definitive endoderm (EOMES, SOX17 and FOXA2) and primitive streak markers (MIXL1, GSC, LHX1), and they were predominantly mapped to anterior primitive streak in the mouse atlas (Figures 4B, 4C and S6C). We identified an HIO epithelial trajectory (c0/2/6/7/11/12/14) marked by the co-expression of CDX2 and CDH1, and predominantly mapped to the mouse gut epithelium (Figures 4C and S6B). We also identified a mesenchymal trajectory (c1/3/5/8/10) that showed the highest similarity to mesenchyme/mesoderm populations in the mouse atlas. We noted that while cluster 9 was CDX2+/CDH1+ and showed comparable similarity to gut epithelium and surface ectoderm, it also exhibited mesoderm features (HAND1+/FOXF1+) suggesting bipotency (Figures 4C, S6B-S6D). In addition, we observed low-abundance neural-like (c15/18), endothelial-like (c16), and gut epithelium-surface ectoderm-like cells (c11) (Figures 4C, S6B and S6D). These cell types decreased after 14 days in culture, and 4-week HIOs were predominantly composed of epithelial and mesenchymal cell populations (Figure 4C).

**Figure 4.**
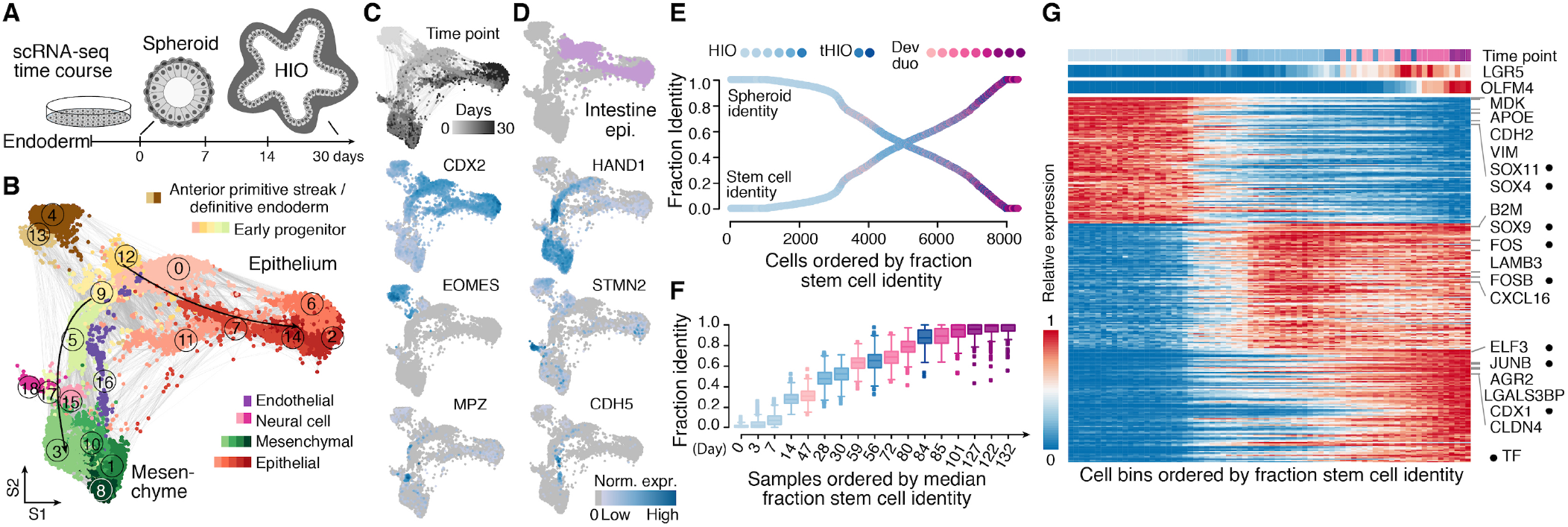
HIOs enable tracking of molecular transitions during small intestinal stem cell development and maturation. A) Schematic of scRNA-seq experiments performed over a time course of HIO development (endoderm, d0-spheroid, d3, d7, d14, d28/30 HIO) to track the progression of epithelium and mesenchyme cell states. B) SPRING embedding colored by cell class and shaded by cluster assignment. Note red and green shades represent epithelial and mesenchyme cells, respectively. C) HIO time course UMAP embedding colored by time point or marker gene expression. D) Light purple cells show HIO epithelial cells that have acquired intestinal identity. E) The fraction identity to spheroid or day 132 developing duodenum stem cell average transcriptome was calculated for each HIO, tHIO, and primary developing duodenum stem cells in all time points. Cells were ordered by increasing fraction stem cell identity and colored by time point and tissue source. F) Distributions of fraction stem cell identity estimated by quadratic programming for each sample ordered by median. Age of tHIO = in vitro period (4 weeks) + post-transplantation period, so as to be comparable with age of HIO, i.e. 4- and 8-week tHIO are day 56 and day 84. Boxplots colored by time point and tissue source. G) Heatmap shows molecular transitions during the acquisition of ISC identity. Cells are ordered according to fraction stem cell identity. Every 50 cells were grouped into a bin, and top sidebars show the most frequent time point, average LGR5, and OLFM4 expression within each cell bin. Selected transcription factors (TFs, circles), ligand and receptor genes are marked next to the heatmap.

To reconstruct an ISC maturation roadmap, we first selected intestine-committed epithelial cells from the in vitro HIO time course (Figures 4D and S6E) (based on CDX2 expression or mapping results to the developing intestine reference, see also STAR methods), and then combined these cells with stem cells from tHIOs and the developing duodenum. We used quadratic programming [44] to decompose the transcriptome of each cell into an early intestinal progenitor (day 0-spheroid) or developing ISC (day 132) component defined in our data, and ordered cells based on increasing developing ISC identity (Figure 4E). We found the fractional ISC identity increased gradually over time, and noted a correspondence between organoid and developing duodenum time points in terms of stem cell maturation. In this case, data from day 47, 59, and 85 tissues showed the most similar ISC identity to day 14 HIO, 4- and 8-week tHIO, respectively (Figure 4F). We searched for genes showing variable expression along this maturation trajectory, and identified genes that are downregulated upon ISC specification (around 14 days) and genes that are upregulated during early (around 4-week HIO) and late phases (emergence of LGR5+ cells) of stem cell maturation (Figure 4G; Table S4). GO enrichment analysis (Table S4) revealed that genes involved in cell adhesion (e.g. VIM, CDH2, VCAN, FN1), and developmental processes (e.g. SOX11, SOX4, CALB1) are downregulated early on in the stem cell trajectory. A diverse set of genes involved in transcriptional regulation (SOX9, FOS, FOSB), oxygen and nitric oxide sensing (NOSTRIN, MTDH, CYBA), focal adhesions and signaling (ROCK2, MAPK3, LGALS1), and apoptosis inhibition (SQSTM1, TPT1, FABP1, PLAC8, CIB1) are upregulated at early stages of ISC development. At later stages, genes involved in microvillus organization (KLF5, VIL1, USH1C, CDHR5), metal ion homeostasis (MT2A, MT1E, FBP1, SLC40A1), defense response (NFKBIZ, ELF3, ASS1, CLU, TNFRSF14), and metabolism (ASS1, CPS1, GLS) are upregulated. This data shows that integrating HIO and primary tissue data enables the temporal reconstruction of ISC differentiation from early progenitor stages, which is difficult to access through primary data, to a mature stage.

### Developing human duodenum mesenchyme is diverse and recapitulated in tHIOs

Mesenchymal-epithelial signaling is vital during intestinal epithelial specification and stem cell maturation during development, and to maintain crypt-villus homeostasis in adulthood [30, 45–48]. We observed multiple mesenchymal cell clusters in the HIO time course and tHIO datasets, however, it was unclear how these cells relate to primary counterparts. To this end, we analyzed cellular heterogeneity in the developing human duodenum mesenchyme. We identified known mesenchymal subpopulations, such as vascular smooth muscle cells (VSMC, c2, DES+/RGS5+) and interstitial cells of Cajal (ICC, c4, ANO1+/KIT+) (Figures 5A, 5B and S7B; Table S2), as well as novel subpopulations, such as CRABP1+ (c3, CRABP1+/SFRP1+) and GDF10+ (c8, GDF10+/C1QTNF3+) (Figures 5A, 5B and S7B; Table S2). We found that mesenchymal subtype composition changed across the time span of developing duodenum samples used in this study (Figure S7A). While mesenchymal subtype CRABP1+ (c3) and ICC (c4) were predominantly observed in day 47, GDF10+ (c8) and SMC (c7) expanded at around day 80 and dominated the day 132 mesenchymal populations (Figure S7A). Around day 80 also marks the proportional decrease of proliferative mesenchyme (c10, MKI67+/SFRP1-, Figure S7A). Through comparison to the multi-organ atlas, we further found that certain mesenchyme subtypes observed in developing duodenum were specific to GI tissues, such as F3+ subepithelial mesenchyme (c15, subepithelial, NRG1+/F3+) and GPX3+ villus core (c13, GPX3+/EDIL3+). We also observed mesenchyme subtypes that were specific to non-GI tissues, such as chondrocyte-like mesenchyme (c21, SOX9+/COL9A3+) in the lung (Figure S8A-C; Table S2). We identified transcription factors (TFs) showing enriched expression levels in mesenchymal subtypes (Figure S8D), with SOX6 being an example of a TF enriched in intestinal subepithelial mesenchyme and its tHIO counterpart (Figure S8E).

**Figure 5.**
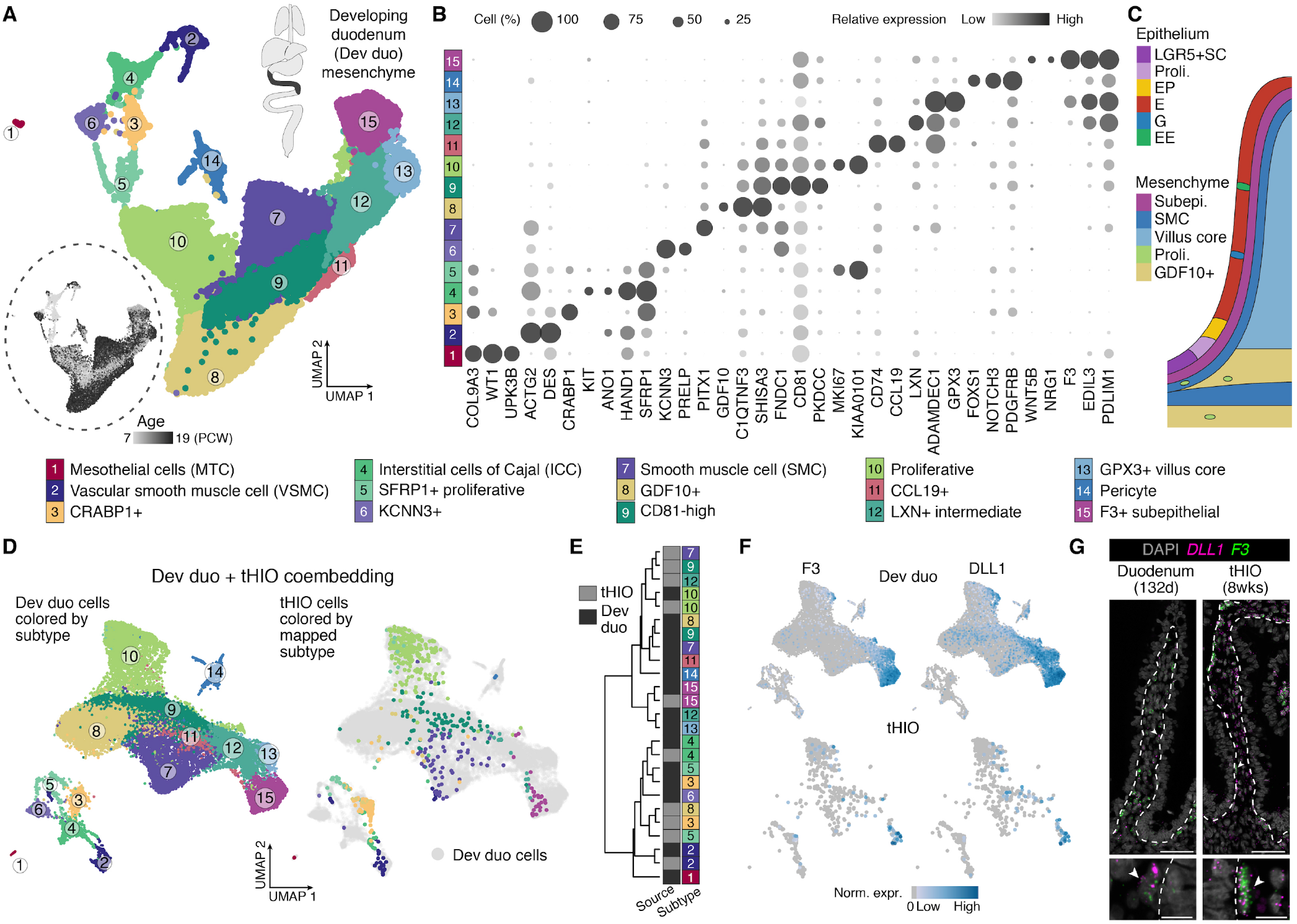
Developing human duodenum mesenchyme is diverse and cell states are recapitulated in tHIOs. A) UMAP embedding of mesenchymal cells from developing human early small intestine and duodenum, with cells colored by cell types or age (bottom left). Cell type annotations are shown below the plot. B) Dot plot shows the average gene expression levels (color) and expressed proportion (size) of cell type enriched genes in each subtype. C) Schematic of epithelial and mesenchymal subtype pseudospatial distribution on day 132 developing duodenum reconstructed with novoSpaRc based on in situ hybridization data [30, 52]. Enterocyte, E; Enteroendocrine, EE; MUC2+ goblet, G; Enterocyte precursor, EP; Intestinal stem cell, SC; Subepithelial mesenchyme, Subepi; TAGLN+ smooth muscle cell populations, SMC; Proliferative mesenchyme, Proli; GDF10+ mesenchyme, GDF10+. D) Integrated UMAP embedding of developing duodenum and tHIO mesenchymal cells. Left, the developing duodenum cells on integrated embedding colored by subtype. Right, tHIO cells colored by the most similar developing duodenum subtype; grey cells represent developing duodenum cells. E) Dendrogram showing transcriptome distance between the developing duodenum and tHIO mesenchyme subtypes. Transcriptome distance was calculated by Spearman’s correlation coefficients of top 50 markers of each developing duodenum subtype. tHIO mesenchymal subtype annotation was refined by comparison to the developing duodenum cells. F) Feature plots of subepithelial mesenchyme-enriched genes, F3 (left) and DLL1 (right) in the developing duodenum (top row) and tHIO (bottom row). G) RNA fluorescence in situ hybridization (FISH) of subepithelial mesenchyme enriched genes (DLL1, pink; F3, green) in day 132 developing duodenum (left) and 8-week tHIO (right). Bottom is a magnification of the top image. Scale bars, 50 μm in upper images, 10 μm in magnified view. Arrows point out DLL1+/F3+ mesenchymal cells located in the subepithelial region in the developing duodenum and tHIOs.

Next, we performed receptor-ligand pairing analysis [49, 50] to infer potential interactions between epithelial and mesenchymal subtypes (Table S5). This analysis revealed interacting gene pairs between epithelium and subepithelial mesenchyme that have been previously reported, such as NRG1-ERBB3 [30]. We further constructed a cell type communication network based on the gene pairs with significant interaction between cell types (Figure S7D), and identified epithelial-subepithelial mesenchyme gene pairs that were differential along the crypt-villus axis (Figure S7E). We visualized spatial expression patterns of interacting gene pairs using novoSpaRc [51] based on a reference of RNA in situ hybridization data of duodenum cell type markers [30, 52] (Figures 5C, S7C and S7F), and predicted interaction pairs enriched in the stem cell niche (Figure S7F; Table S5). For example, we found EPHB2 (receptor of Eph/ephrin signaling) and EREG (ligand of EGFR) are enriched in stem cells, and their interaction partners (EFNA5, EGFR respectively) showed slightly enriched expression in subepithelial mesenchyme adjacent to epithelial stem cells (Figure S7F) which was reported to regulate stem cell proliferation and cell positioning in the adult stem cell niche [53]. EREG-EGFR mediated signaling was shown to rescue morphogenesis and suppress apoptosis in intestine organoids derived from Yap-null mice [54]. These data revealed a complex consortium of molecularly distinct duodenal mesenchymal cell fates with features that are specific to intestinal niches.

Furthermore, we compared tHIO mesenchyme subtypes to the developing duodenum mesenchyme reference. Integrated co-embedding of tHIO and the developing duodenum mesenchyme revealed similarities of mesenchymal cells, suggesting tHIO mesenchyme subtypes recapitulate transcriptome features of the primary counterparts (Figures 5D and S7G). Hierarchical clustering based on expression patterns of cell type markers identified in the primary reference further shows that VSMC, ICC, subepithelial and proliferative mesenchyme in tHIO and the developing intestine co-cluster (Figures 5E and S7G). Next, we used in situ hybridization to validate localization of the F3+/DLL1+ mesenchyme subtype (c15 in the developing duodenum mesenchyme reference) at subepithelial mesenchyme locations of both tHIO and the developing duodenum (Figures 5F and 5G). Moreover, we analyzed the mesenchymal cells and early progenitors of in vitro HIOs and identified putative precursors to smooth muscle, subepithelial/intermediate/villus core, and CRABP1+ mesenchymal cells that were observed in the developing duodenum (Figures S9A-S9F). Together, this data shows that mesenchymal cells fates emerge in HIOs that match to the developing small intestine counterparts.

### CDX2 deletion leads to loss of intestinal cell fate and gain of foregut features in both epithelium and mesenchyme

We next utilized HIOs to understand the regulatory mechanisms that lead to the specification of intestinal epithelium and mesenchyme cell type identities. We used the HIO time course scRNA-seq data to construct a transcription factor (TF) network based on expression correlation (Figure 6A) and identified TFs that distinguish progenitor, epithelial, and mesenchymal cell states (Figure 6B). This analysis revealed differentially expressed TF modules that might drive the diversification of epithelium and mesenchyme from early HIO time point progenitors. Interestingly, we find that CDX2, a known master regulator of intestine epithelium [11, 55–59] and critical regulator of mesoderm cell fate specification [60, 61] is positioned in the graph with genes that have correlated expression in early progenitors (Figures 6A and 6B). CDX2 is highly expressed in HIO epithelial cells, and its expression is maintained in the developing small intestine epithelium into adulthood. Interestingly, CDX2 is also expressed in a subset of early mesenchymal progenitors in the HIO (c5, c9), as well as mesoderm/mesenchyme populations in the mouse gastrulation atlas (Figures 6C and S10A-C).

**Figure 6.**
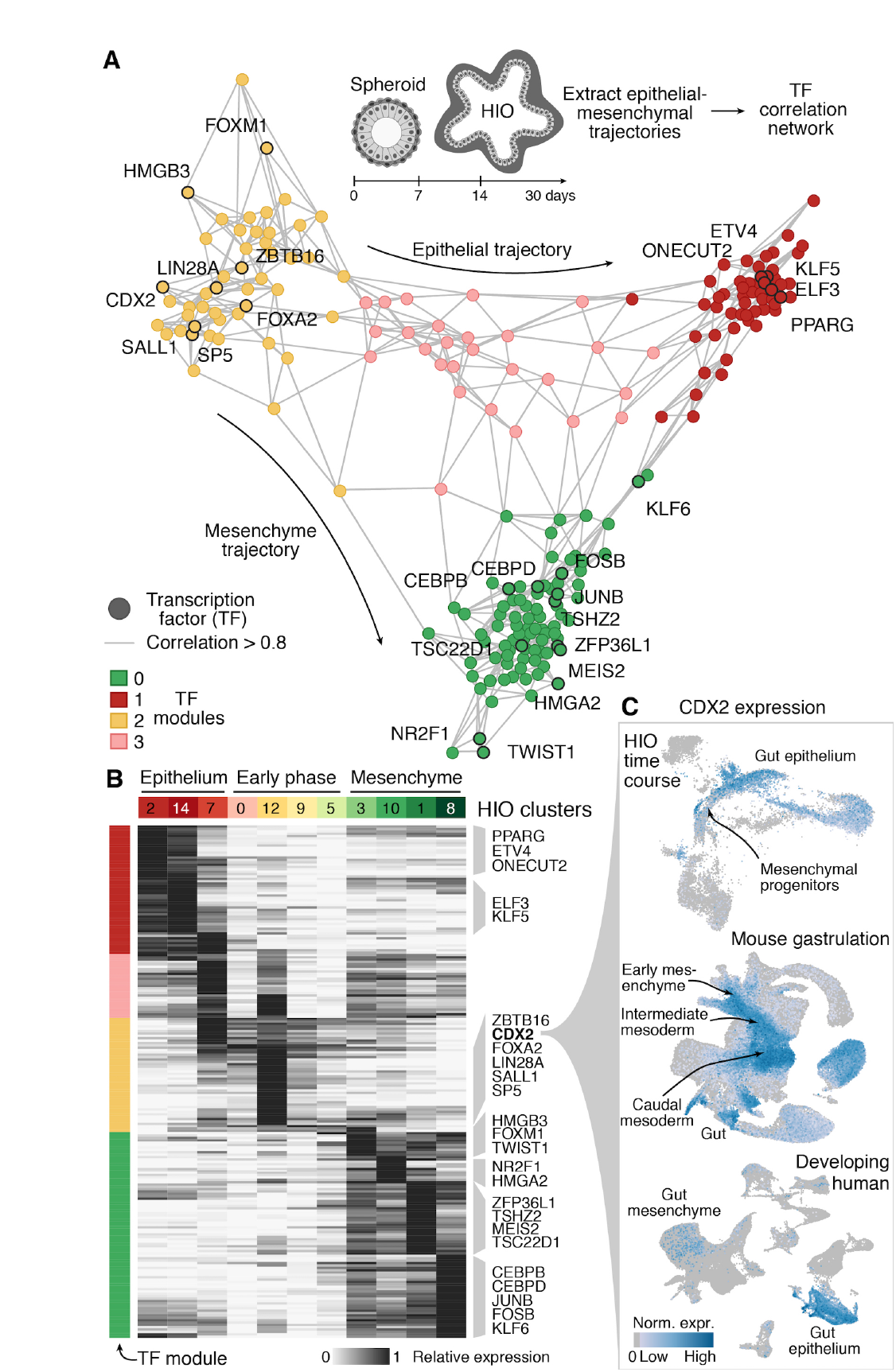
Transcription factors correlate with intestine epithelial and mesenchyme specification. A) Transcription factor correlation network during HIO epithelial and mesenchymal development. Shown are nodes (TFs) with no less than ten edges, colored by expression module, with each edge representing a high correlation (>0.8) between connected TFs. The top HIO cluster-enriched TFs are labeled. B) Heatmap showing the expression of TFs in each module across early phase, epithelium and mesenchyme HIO clusters, with TFs highlighted from the network graph. HIO clusters are the same as those shown in Figure 4B. C) Expression of intestinal master regulator CDX2 in HIOs, mouse gastrulation, and the developing endoderm cell atlases.

To test the requirement of CDX2 to specify both gut epithelium and mesenchyme fates, we analyzed HIOs generated from hPSCs harboring homozygous CDX2 knockout (CDX2-KO) and a corresponding control line [11] (Figure 7A). Immunohistochemistry and scRNA-seq data supported that CDX2 expression was absent in CDX2-KO organoids (Figures 7B and 7C). Consistent with previous reports [11, 59, 62], CDX2-KO organoids exhibited predominantly cystic morphology of folded and glandular epithelium (Figures 7B and S11A), and were positive for canonical foregut markers SOX2 and MUC5AC (Figure 7B). We compared scRNA-seq data from control and CDX2-KO organoids, and CSS integration revealed cell type composition differences between the two conditions, with multiple epithelial cell populations being specific to CDX2-KO organoids, such as MUC5AC+ (c7, MUC5AC+/CLDN18+), MUC16+ (c11, MUC16+/AQP3+), and SPP1+ (c10, SPP1+/HABP2+) cells (Figures 7B-7D and S11B; Table S2). Comparing epithelial cells in CDX2-KO organoids to the developing human reference atlas revealed that these cells were predominantly mapped to the stomach epithelium (Figure 7E). Together, this data indicates a loss of intestine and expansion of stomach identities in CDX2-KO organoid epithelium.

**Figure 7.**
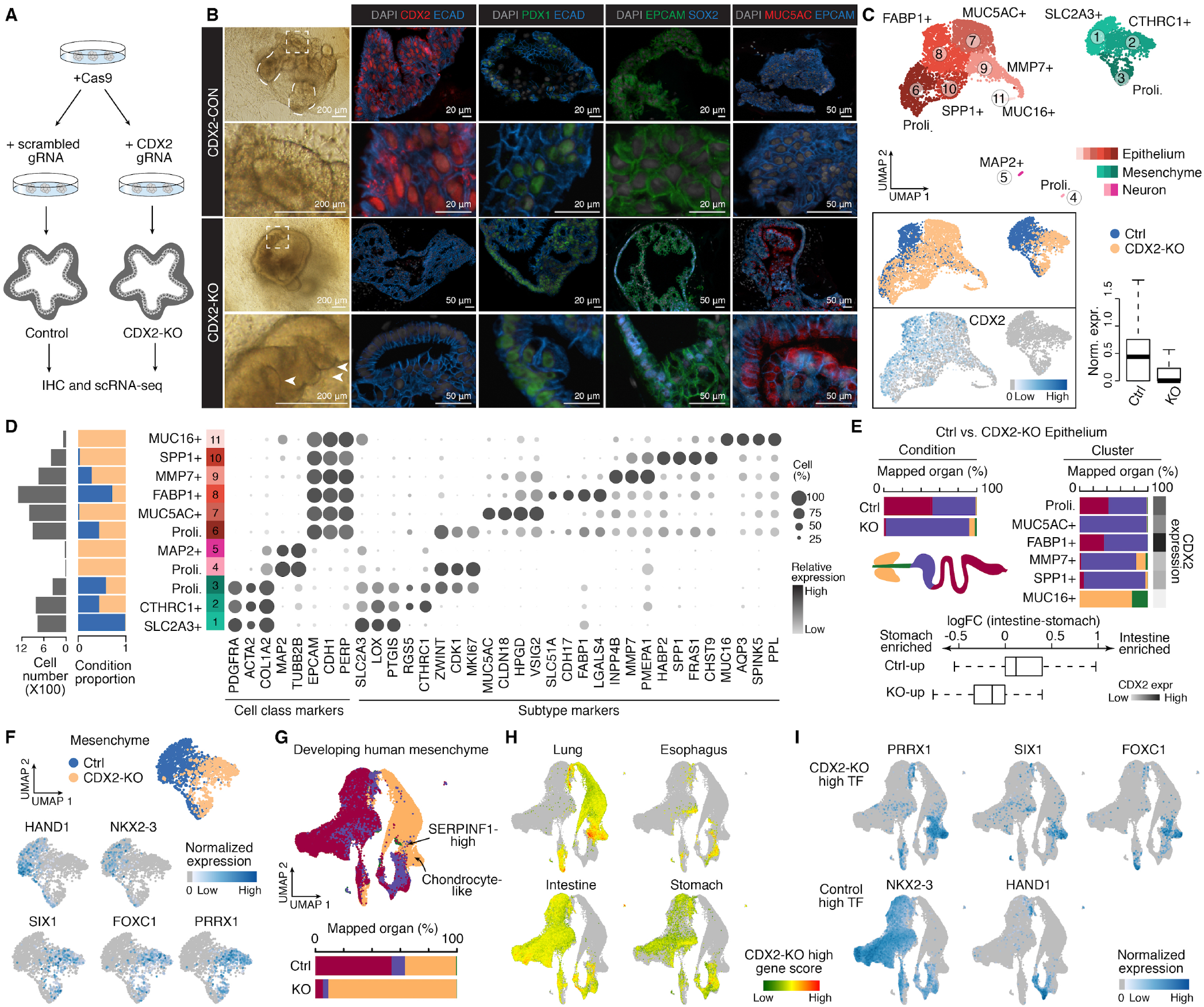
CDX2 deletion leads to loss of intestinal cell fate and gain of foregut features in both epithelium and mesenchyme. A) Schematic of CDX2-knockout (KO) experiment [11]. B) Left, bright field (BF) images show the morphology of control (top) and KO (bottom) organoids. Dashed curves of control organoid image show protrusions of crypt-like structure. Dashed square and magnified control organoid image show typical intestine epithelium structure. Dashed square and magnified KO organoid image show folded glandular epithelial organization (white arrow). Right, immunofluorescence (IF) stainings of organ enriched epithelial markers (intestine, CDX2; proximal small intestine, PDX1; foregut, SOX2, MUC5AC) in control (top) and KO (bottom) organoids. Scale bars, 200 μm in BF images; 20 μm or 50 μm in IF images. C) Integrated UMAP embedding of control and KO organoid cells. Top, cells are colored by major cell class with shade representing cluster assignment (epithelium, red; mesenchyme, green; neuron, pink). Bottom left, cells are colored by condition or CDX2 normalized expression levels. Bottom right, box plot shows CDX2 expression levels in control and KO organoids. Thick lines represent medians. D) Left bar plot shows cell numbers of each cluster. Right stacked bar plot shows proportions of cells from each condition in each cluster. Dot plot shows cluster marker gene expression. E) Top, stacked bar plot shows the proportion of epithelial cells in each condition (left) and each cell cluster (right) mapping to reference atlas organs. The sidebar shows CDX2 expression. Bottom, box plot shows log-transformed fold change of control and KO epithelium differentially expressed genes in the developing intestine versus stomach. F) Feature plots of differentially expressed transcription factors (TF) between mesenchymal cells of control and CDX2-KO organoids. Only mesenchymal cells are shown. G) Top, UMAP embedding of mesenchymal cells subsetted from the developing human reference atlas, with cells colored by organs. Bottom, stacked barplot shows the proportion of mesenchymal cells in each condition mapping to reference atlas organs. Color scheme of organs is the same as panel E. H-I) Mesenchyme UMAP colored by cumulative z-transformed expression levels of top 20 positive markers of CDX2-KO mesenchyme in each developing organ (H), or expression of differentially expressed TFs between control and CDX2-KO mesenchyme (I).

We next performed differential expression (DE) analysis between control and CDX2-KO mesenchyme (Figures S11C and S11D; Table S6), and examined the expression patterns of DE genes in the mouse gastrulation atlas and HIO time course (Figures S11E-S11I), as well as in the developing human reference. We noted that genes with higher expression in CDX2-KO mesenchyme showed enriched expression in the lung subtypes (Figures 7F-7I). For example, TFs highly enriched in CDX2-KO (e.g FOXC1, PRRX1, SOX9 and SIX1) shared enriched expression in the lung chondrocyte-like mesenchyme (c21, Figure S8A) and SERPINF1-high (c20, Figure S8A) mesenchyme (Figures 7F and 7G). SIX1 and PRRX1 are required for morphogenesis [63, 64] and vascular development in the developing lung [65, 66], respectively. Meanwhile, we observed a loss of expression of the ubiquitous intestine mesenchyme marker TF NKX2-3 [67, 68] in CDX2-KO mesenchyme (Figures 7F and 7I). We further examined CDX2-KO induced alteration of organ identity in mesenchymal cells according to a k-nearest neighbour (k = 20) search in the developing endoderm cell atlas. We found that CDX2-KO mesenchyme tends to be mapped to the developing lung over the intestine as compared to control mesenchyme (Figure 7G, Fisher’s exact test, nominal P < 0.0001, odds ratio = 25.23). These data suggest CDX2-KO induced the loss of intestine identity and gain of certain lung mesenchymal features in the HIOs. Taken together, we show that CDX2 is critical for both epithelial and mesenchymal cell fate acquisition during human early intestine development, and provide a groundwork for further interrogation of mechanisms controlling epithelial and mesenchymal specification and interactions in the human intestine.

## Discussions

Here we set out to understand how epithelial and mesenchymal cells develop within the HIOs using single-cell transcriptomics. By generating a reference atlas from diverse endoderm-derived human organs, and leveraging comparisons with the mouse gastrulation atlas, we were able to rigorously assess the cell composition of HIOs. We identify off-target, non-intestinal cell types that are present in the culture system, which could impact interpretations from experiments that lack single-cell resolution. We find that the majority of HIO cells, both in vitro HIOs and transplanted HIOs, are specified to intestinal fates, and the transplanted HIOs have a remarkable resemblance to the developing human intestine on the morphological and molecular level. Furthermore, we find each of the major epithelial cell types in the transplanted organoids is more similar to the developing counterparts than adult, including stem cells, differentiated enterocytes, enteroendocrine, and goblet cells. This feature provides opportunities to understand the mechanisms that underlie the maturation of human intestinal cell types.

We also provided an analysis of the molecular transitions that take place in ISCs from early stages of human development into adulthood, and these findings highlight intriguing transcription factors, signaling pathways, and metabolic features. These features could be targeted to enhance HIO stem cell maturation in vitro through transcription factor overexpression [69], CRISPR inhibition/activation [70, 71], or small molecule modulations, each of which could be adapted using high-throughput single-cell genomic [72] or image-based [73] readouts. We also observe that HIOs mature after transplantation into a mouse host; however, we note that there are multiple cell types observed in developing intestine that are underrepresented or absent in HIOs, including endothelial and immune cell populations. Alterations to the HIO culture system that enable co-differentiation or co-culture of endothelium [14], neuron [17], immune or other lineages are interesting approaches to achieve a more complete model of human intestine development, and may allow enhanced maturation of cell types. Furthermore, maturation may also require perfusable vasculature [74, 75], extracellular matrix alterations [10], or other mechanical inductive cues [76]. Achieving mature intestinal tissue from iPSCs in vitro remains a major challenge in the field, and continued benchmarking against multi-organ reference atlases such as the one generated here, will be required to quantify the precision of cell state specification. Nonetheless, the molecular, cellular, and morphological similarities between tHIO and the developing duodenum epithelial and mesenchymal cell types are striking and present extraordinary opportunities to use tHIOs as a predictive model system.

Currently, there is very little known about the genetic mechanisms that govern human intestinal mesenchyme development. Here, we provide a single-cell transcriptome reference of developing duodenum mesenchyme, and show that tHIOs develop remarkably similar counterparts, thereby providing a system to explore intestinal mesenchyme in controlled culture environments. It has been shown that subepithelial mesenchyme promotes ISC maturation through epithelial-mesenchymal interactions [77–79]; however, the signaling and transcriptional networks that regulate human intestinal sub-epithelial mesenchyme differentiation are only starting to be understood from single-cell transcriptomics [30, 31]. Notably, major duodenum mesenchyme clusters have tHIO counterparts, and we identify early precursor states within the in vitro HIO. This provides a system to understand the gene regulatory logic controlling intestinal mesenchyme subtype specification.

Our findings that CDX2 is critical for human intestinal epithelial differentiation is consistent with previous reports in mice. Cdx2 deletion in mice results in a homeotic transformation of the intestine into an esophagus-like tube [59] or ectopic expression of stomach markers [57], depending on the developmental timing of Cdx2 loss. Consistent with previous animal studies showing Cdx2 plays a role in mesoderm posteriorization [60, 61], we show that during HIO development, loss of CDX2 leads to anteriorization of mesenchymal fates such that CDX2-KO mesenchyme loses intestine identity and acquires lung-associated mesenchyme identity. Our analyses of HIO developmental trajectories, suggests that CDX2 is a transient patterning regulator during early phases of gut-associated mesenchyme development in humans, and CDX2 expression is not required to maintain the intestine identity in mesenchyme. In addition, the difference of CDX2-KO induced regional identity alteration in epithelium and mesenchyme indicates that regional identity of mesoderm is not dependent on epithelial identity. In line with that, a previous study [59] showed that epithelial-specific deletion of Cdx2 in mice resulted in mispatterning of the epithelium, whereas the mesenchymal Hox gene expression patterns were largely intact, indicating that the regional identity of the mesoderm is maintained in epithelial-CDX2 knockout intestines. To our knowledge, our study is the first example showing that iPSC-derived human organoids reveal the consequence of a genetic perturbation on regional identity and patterning.

Altogether, our data and analyses provide highly-resolved evidence that HIOs can be predictive of human intestinal epithelial and mesenchyme development, and that perturbations in HIOs can reveal developmental mechanisms in human tissues. Furthermore, we highlight genes that are associated with neonatal and pediatric digestive disorders and find that epithelial, as well as mesenchymal cell states, are implicated in these disorders (Figure S12; Table S7). Because of the cellular heterogeneity and presence of non-intestinal cell types observed in HIOs, it will be important to use single-cell genomic analyses of patient-derived organoids and other HIO models of disease to appropriately resolve associated phenotypes. Lastly, our data points to the value of high-dimensional human reference cell atlases across development, adulthood and disease, to comprehensively benchmark engineered human tissue culture systems, and such comparisons could be used to improve organoids for improved modeling of development and disease.

## ACKNOWLEDGEMENTS

We thank the Camp and Treutlein labs for helpful discussions. We thank Judy Opp and the University of Michigan Advanced Genomics Core for their expertise in operating the 10X Chromium single-cell capture platform and sequencing. We would also like to thank the University of Washington Laboratory of Developmental Biology staff for their support on this project. JGC, JRS and BT are supported by grant number CZF2019-002440 from the Chan Zuckerberg Initiative DAF, an advised fund of Silicon Valley Community Foundation. JGC is supported by the European Research Council (Anthropoid-803441) and the Swiss National Science Foundation (Project Grant-310030_84795). BT is supported by the European Research Council (Organomics-758877, Braintime-874606), the Swiss National Science Foundation (Project Grant-310030_192604) and the National Center of Competence in Research Molecular Systems Engineering. JRS is supported by the Intestinal Stem Cell Consortium (U01DK103141), a collaborative research project funded by the National Institute of Diabetes and Digestive and Kidney Diseases (NIDDK) and the National Institute of Allergy and Infectious Diseases (NIAID). JRS is also supported by the National Heart, Lung, and Blood Institute (NHLBI - R01HL119215), by the NIAID Novel Alternative Model Systems for Enteric Diseases (NAMSED) consortium (U19AI116482). IAG and the University of Washington Laboratory of Developmental Biology were supported by NIH award number 5R24HD000836 from the Eunice Kennedy Shriver National Institute of Child Health and Human Development (NICHD). EMH was supported by the Training in Basic and Translational Digestive Sciences Training Grant (NIH-NIDDK 5T32DK094775), the Cellular Biotechnology Training Program Training Grant (NIH-NIGMS 2T32GM008353), and the Ruth L. Kirschstein Predoctoral Individual National Research Service Award (NIH-NHLBI F31HL146162). MC was supported by the Training Program in Organogenesis (NIH-NICHD T32 HD007505). Additional support was provided by the University of Michigan Center for Gastrointestinal Research (UMCGR) (NIDDK 5P30DK034933).

## AUTHOR CONTRIBUTIONS

UK, EMH and YHT grew organoids. YHT and MC transplanted organoids into mice hosts. UK, AW, YHT, MC and EMH collected, dissociated and submitted tissues for scRNA-seq. QY and ZH performed scRNA-seq data analysis. JHW and QY maintained the scRNA-seq data. YHT and UK performed immunofluorescence stainings. EMH and CC performed multiplexed fluorescence in situ hybridization and imaging. YHT performed chromogenic in situ hybridization. IAG and PDRH provided critical material for this work. QY, UK, EMH, BT, JRS and JGC designed the study and wrote the manuscript. All authors read and approved the manuscript.

## DATA AVAILABILITY

Submission of raw mRNA sequencing data is in process. Processed sequencing data and reference patterns for novoSpaRc have been deposited in Mendeley Data with http://dx.doi.org/10.17632/x53tts3zfr.1 and is currently in private status. Please contact authors for inquiries.

## CODE AVAILABILITY

All code used for single-cell analysis and data presentation is available via GitHub at https://github.com/camplab-iob/humanintestineorganoids.

## STAR Methods

### Human subjects

Normal, de-identified human adult intestinal biopsies were collected from a 45-year-old male and a 50-year-old female with approval from the University of Michigan IRB. Biopsy specimens were stored on ice in sterile saline prior to single-cell dissociation. Normal, de-identified developing human tissues were obtained from the University of Washington Laboratory of Developmental Biology, and all work was approved by the University of Washington and the University of Michigan IRB. Tissue was shipped overnight in Belzer-UW Cold Storage Solution (ThermoFisher, NC0952695) with cold packs. A list of tissue specimens can be found in Table S1.

### Generation of human intestinal organoids

Human ESC line H9 (NIH registry 0062) was obtained from the WiCell Research Institute and CDX2-control and CDX2-knockout lines used in this study were generated on the H9-background [11]. All experiments using human ES cells were approved by the University of Michigan Human Pluripotent Stem Cell Research Oversight Committee. hESCs differentiated into human intestinal organoids (HIOs) based on the previously described protocol [1, 6]. Briefly, hESCs were patterned into definitive endoderm (DE) by Activin A (100 ng/ml) in RPMI-1640 media for 3 days with increasing concentrations of HyClone FBS respectively (0%, 0.2%, 2%). Midgut/hindgut patterning was carried out in 2% HyClone FBS containing RPMI-1640 supplemented with FGF4 (500 ng/ml) [81] and CHIR99021 (2 μm). Media was changed daily, and spheroids were collected at the end of 5 days of midgut/hindgut patterning. Next, spheroids were embedded in Matrigel (8 mg/ml, Corning, 354234) and incubated in ENR media (mini gut basal media, supplemented with EGF (RD Systems, 236-EG-01M, 100 ng/mL,), Noggin-Fc (100ng/mL) (purified from conditioned media [82]), and R-Spondin2 (5% conditioned medium [83]). The media was changed every 4-5 days. The tHIOs processed for FISH were grown in ENR media for the first 72 hours days to pattern duodenal identity, after which media was changed to only contain EGF (100ng/ml) in mini gut basal media. The mini gut basal media is composed of the following components: Advanced DMEM: F12 (Life Technologies, 12634), 1x B27 supplement (Life Technologies, 17504044), 2 mM L-Glutamine (Life Technologies, 25030), 15 mM HEPES (Life Technologies, 15630080). All media used in the differentiation process contain 1x penicillin and streptomycin (Life Technologies, 15140).

### Human intestinal organoid (HIO) transplantation

The University of Michigan Animal Care and Use Committees approved all animal research performed in this study. Firstly, 4-week HIOs were mechanically dissociated from Matrigel. Then, HIOs were implanted under the kidney capsules of immunocompromised NOD-SCID IL2Rg-null (NSG) mice (Jackson Laboratory strain no. 0005557) as previously described [7, 8]. In summary, mice were anesthetized using 2% isoflurane. A left-flank incision was used to expose the kidney after shaving and sterilization with isopropyl alcohol. HIOs were implanted beneath mouse kidney capsules using forceps. Prior to closure, an intraperitoneal flush of Zosyn (100 mg/kg; Pfizer) was administered. Mice were euthanized for retrieval of tHIOs after 4 and 8 weeks for scRNA-seq and 10 weeks for stainings. Results shown are representative of two experiments performed with a total of 8 mice.

### Chromogenic In Situ Hybridization and Multiplex Fluorescent In Situ Hybridization (FISH)

Paraffin blocks were sectioned to generate 5 μm-thick sections. Sections were mounted to SuperFrost Plus Slides (Thermo Scientific, 10149870) and used within one week for optimal results. The assay was carried out in RNase-free conditions by treating all materials with RNase-away (Molecular Bioproducts Inc., 7005-11) prior to use. Slides were stored at room temperature in a sealed slide box with silica desiccant packets. Slides were baked for 1 hour in a 60°C dry oven a day prior to starting the procedure. Chromogenic in situ hybridization (ISH) and multiplex fluorescent in situ hybridization (FISH) protocols were performed according to the manufacturer’s instructions (ISH–ACD; RNAscope 2.5 Assay BROWN 322310-QCK Rev B; FISH-ACD; RNAscope Multiplex Fluorescent v2 manual protocol, 323100-USM), under standard antigen retrieval conditions (15 min) and optimized protease treatment conditions for each tissue (intestine–30 min, tHIO–20 min). FISH samples were imaged using Nikon A1 confocal and images were assembled using Photoshop CC. Imaging parameters were kept consistent for all images within the same experiment and any post-imaging manipulations (i.e. brightness, contrast, LUTs) were performed equally on all images from a single experiment.

### Tissue stainings

Immunofluorescence stainings were conducted as previously described [84]. Briefly, tissues were fixed overnight in either 4% PFA or 10% neutral buffered formalin (NBF) for approximately 24 hours at room temperature. The following day, tissues were washed in UltraPure Distilled Water (Invitrogen, 10977-015) for 3 changes for a total of 2 hours. Tissue was gradually dehydrated in a methanol series (25%, 50%, 75%, 100%). Tissue was stored long-term in 100% Methanol at 4°C. Prior to paraffin processing, tissue was equilibrated in 100% ethanol for an hour followed by 70% ethanol. Tissue was paraffin perfused using an automated tissue processor (Leica ASP300) with 1-hour solution changes overnight. Paraffin processed tissue was embedded and 5-7μm sections were cut.

Paraffin sections were first deparaffinized in Histoclear and re-hydrated. Antigen retrieval was performed by steaming slides in a sodium citrate buffer for 20 minutes. Slides underwent a blocking step using the appropriate serum (5% serum in PBS + 0.5% Triton-X) for 1 hour at room temperature. Primary antibodies were diluted in blocking solution (1:500) and slides were incubated with antibodies overnight at 4°C. The following day, slides were washed and incubated with appropriate secondary antibodies (1:500) diluted in a blocking buffer for 1 hour at room temperature together with DAPI staining (1μg/mL). Slides were washed and mounted using Prolong Gold (Thermo Fisher, P10144). A list of can be found in the Key Resources Table.

HE staining was performed with Harris Modified Hematoxylin (Fisher Scientific, SH26-500D) and Shandon Eosin Y (Thermo Scientific, 6766007) based on the manufacturer’s instructions.

### Single-cell dissociation and RNA sequencing library preparation of the developing human tissues

Developing human tissue and organoid dissociations for scRNA-seq were performed according to a previously published study [85]. Initially, all tubes and pipette tips were pre-washed with 1% BSA in HBSS to prevent adhesion of cells to the plastic. Organoids were mechanically removed from Matrigel droplets. Next, organoids and primary tissues minced into small fragments, using a scalpel in a petri dish filled with ice-cold 1X HBSS. Then, tissue and organoid fragments were transferred into a 15 mL conical tube. Enzymatic dissociation started in the Neural Tissue Dissociation Kit (Miltenyi Biotec, 130-092-628), and all incubation and centrifugation steps were carried out in a refrigerated centrifuge pre-chilled to 10°C for the developing tissue samples, whereas, 37°C for organoid samples unless otherwise stated. Based on the manufacturer’s instructions, the fragments of tissue were treated for 15 minutes with Mix 1. Mix 2 was added to the digestion, and tissue was incubated for 10-minute increments until digestion was complete. After each 10 minute incubation, tissue was agitated using a P1000, and tissue digestion was visually assessed under a stereomicroscope. This process continued until the tissue was fully digested. Cells were filtered through a 70 μm filter coated with 1% BSA in 1X HBSS, spun down at 500g for 5 minutes at 10°C, and resuspended in 500μl 1X HBSS (with Mg2+, Ca2+). 1 mL red blood cell lysis buffer (Roche, 11814389001) was then added to the tube and the cell mixture was placed on a rocker for 15 minutes at 4°C only for the primary tissue samples. Cells were spun down (500g for 5 minutes at 10°C) and washed twice by suspension in 2 mL of HBSS +1% BSA followed by centrifugation. Two exceptions were definitive endoderm and spheroid (d0 HIOs) samples were dissociated using TrypLE™ Express (Gibco, 12605-010) at 37°C. Cells were counted using a hemocytometer and kept on ice. Single-cell droplets were immediately prepared on the 10x Chromium according to manufacturer instructions at the University of Michigan The Advanced Genomics Core, with a target of capturing 5,000-10,000 cells. Single-cell libraries were prepared using the 10x Chromium Single Cell 3 v2 and Next GEM Single Cell 3’ v3.1 kits according to manufacturer instructions.

### Alignment of single-cell transcriptomes

We used Cell Ranger (10x Genomics) to demultiplex base call files to FASTQ files and align reads. Default alignment parameters were used to align reads to the human reference genome provided by Cell Ranger. In vitro organoid time course data and human primary tissue data were mapped to hg19. CDX2 control and knockout line-derived organoid data was mapped to hg38. Transplanted organoid data were mapped to the human-mouse dual genome (hg19 and mm10).

### Analysis of single-cell RNA-seq data

Raw scRNA-seq data of developing human duodenum and early small intestine, lung and subset of in vitro human intestinal organoid data (days 0 and 3) were retrieved from ArrayExpress database (E-MTAB-8221) [14, 30, 38].

In vitro HIO time course, CDX2 knockout and control HIOs, tHIO, developing multi-organ atlas and adult duodenum data were analyzed individually. Seurat (v3.1) package [86] was applied to the scRNAseq data for preprocessing. Generally, cells with less than 1,000 or more than 20,000 detected genes, as well as those with mitochondrial transcript proportion higher than 10% (all except adult duodenum samples) or 50% (adult duodenum samples) were excluded. For the developing human multi-organ cell atlas data, ribosomal genes, mitochondrial genes and genes located on sex chromosomes were removed. After log-normalization, 2,000 or 3,000 highly variable genes were identified using the default vst method. The normalized expression levels were then z-transformed, followed by principal component analysis (PCA) for dimension reduction. Uniform manifold approximation and projection (UMAP) was applied to the top 20 principal components (PCs) for visualization.

To integrate data of different samples, Cluster Similarity Spectrum (CSS) [39] was calculated as described. In brief, cells from each organoid were subsetted, and Louvain clustering (with resolution 0.6), implemented in Seurat, was applied based on the pre-calculated top 20 PCs. Average expression of the pre-defined highly variable genes was calculated for each cluster in each organoid. Afterwards, Spearman’s correlation coefficient was calculated between every cell and every cluster in samples. For each cell, its correlations with different clusters of each sample were z-transformed. Its z-transformed similarities to clusters of different samples were then concatenated as the final CSS representation. All data presented in this manuscript were after CSS integration, except for integrated UMAP embedding of tHIO and 4-week in vitro HIO cells. UMAP and Louvain clustering were applied to the CSS representation.

To integrate tHIO and 4-week in vitro HIO cells, we used the RunHarmony function implemented in the harmony R package [87], with default settings except for using top 20 PCs. UMAP was applied to the harmony representation.

For time course in vitro HIO dataset, its SPRING-based cell embedding for visualization was generated as previously described [88]. In brief, single cells from the same cluster of the same sample with similar transcriptome were grouped into pseudo-cells. Correlation distance between CSS of each pair of pseudo-cells was calculated and a k-nearest neighbour (kNN) network (k = 20) was then calculated with the constraint to only consider pseudo-cells from the same or nearby stages when screening for nearest neighbours. The kNN network was visualized using SPRING [43]. Coordinates of single cells were predicted from pseudo-cells based on CSS using support vector regression (SVR) implemented in the e1071 package. Each SVR model was trained for one dimension of coordinates. Such coordinates were further refined by pushing each cell to its nearest pseudo-cell with the smallest correlation distance of CSS to be 80% closer.

Cluster annotation was done based on expression patterns of canonical cell type markers found in literature, together with cluster markers identified using the presto package. For analysis of tHIO, developing intestine epithelium, duodenum epithelium, duodenum mesenchyme, and adult duodenum, clusters with similar transcriptome and shared annotation were merged. Clusters with marker expression of multiple cell class and without specifically enriched positive markers were considered as doublet clusters, and removed from downstream analysis.

Identification of cluster and cell type markers and differentially expressed genes between categories were identified by calculating area under receiver operator curve (AUC) implemented in the presto package. We used AUC > 0.6, log-transformed expression level fold change > 0.25 as the significance cutoff.

### Quantification of organ specificity

K-nearest neighboring cells (k = 100) were obtained for each cell of the developing human multi-organ atlas by the nn2 function implemented in the RANN package, based on the Euclidean distance in CSS space. For each cell, we calculated the proportions of each organ among its neighboring cells. We also calculated the proportion of each organ in the whole dataset. Our rational is, for a cell, if the proportion of an organ estimated from its neighboring population exceeds that estimated from the whole dataset, and if this organ is different from the organ identity of the cell, i.e. this is a non-self organ, it indicates that this cell tends to share transcriptome similarity with that organ. The larger number of non-self organs fitting this situation indicates this cell intermix with more organs on the transcriptome level, i.e. this cell shows lower organ specificity. Therefore, we could use this number to quantify organ specificity of each cell. The distribution of this number among a group of cells could indicate the organ specificity of that cell population.

### Inference and visualization of inter-cell-class interactions in developing human organs

We retrieved the interacting gene pair annotations from CellPhoneDB [80]. For each developing human organ, genes without any detection in the data set were firstly discarded. Genes with no more than 10 co-expressed genes (Pearson correlation coefficient (PCC) across average expression of cell clusters > 0.5) were also excluded from the following analysis. Next, for each developing human organ, genes were grouped into five groups, according to the cell class (epithelium, mesenchyme, endothelium, immune cells, neural cells) where they reach maximum expression levels. For every two gene groups, each of which represented one cell class, we counted the number of interacting gene pairs where the two genes belong to the two gene groups of interest. A one-sided Fisher’s exact test was then applied to compare the number of interacting gene pairs of one gene group with another one in comparison to the number of intra-group interacting gene pairs, with the sizes of gene groups taken into account. A significant gene group pair suggests the inter-cell-class interactions between the two represented cell classes are more frequent than intra-cell-class interactions, and therefore implies valid interactions or communications between the two cell classes. Nominal P < 0.05 was set as a significance cutoff.

In addition, for each developing human organ, an interacting network of the analyzed genes of each organ was visualized by the igraph R package with force-directed layout in undirected mode, using correlation cutoff (PCC > 0.5)-based adjacent matrix as input, and linked genes with annotated interactions.

### Mapping organoid cells to the developing human multi-organ atlas

First, we calculated the representations of in vitro HIO and tHIO cells which were compatible with the CSS-represented developing human multi-organ atlas. In brief, we calculated the Spearman’s correlation coefficient between query cells and every cluster of each developing human sample using the highly variable genes defined in the developing human atlas. Afterwards, its correlations with different clusters of each developing sample were z-transformed. Its z-transformed similarities to clusters of different samples were then concatenated as the final Reference Similarity Spectrum to the developing human atlas representation (RSS in short), which share the same dimensionality and definitions as the CSS representation used in the atlas.

To allow the projection of the tHIO cells to the UMAP embedding of developing human multi-organ atlas, we first built a UMAP model based on the CSS of the human developing atlas using the ‘umapget_umap_models’ function (with ‘ret_model’ parameter as TRUE) implemented in the uwot package. Based on the RSS of tHIO cells and the pre-trained UMAP model of the developing human endoderm cell atlas data, we used the ‘umap_transform’ function in the uwot package to project tHIO cells to the UMAP embedding of the developing human multi-organ atlas.

To predict the cell type and cell state identities of cells in both in vitro HIO, tHIO samples, as well as the CDX2-KO and control samples, we identified the k-nearest (k = 20) developing human cell neighbors of each in vitro HIO or tHIO cell, by calculating Euclidean distance between RSS of query cells and CSS matrices of the developing human multi-organ cell atlas using the ‘nn2’ function in the RANN package. The most frequent developing human cell identity among the nearest neighbors was defined as the mapped human in vivo identity of each organoid cell.

### Resolving intestinal region specificity of epithelial cell types in developing human and tHIOs

We examined the intestinal region specificity of stem cells, enterocytes, enteroendocrine, goblet and M cells separately. These five cell types were selected as they could be observed in both developing intestine tissues and tHIOs. To avoid confounding effects of individual variations, we only selected developing human specimens with more than three intestine regions collected, i.e. the day 80, 101 and 132 old specimens. For each of the cell types of interest, we first identified genes showing differential expression (DE) levels between small intestine and colon, or between different small intestine regions. Specifically, to identify DE genes between small intestine and colon, we compared each of the collected small intestine tissues to colon for each specimen separately. In each comparison, both groups were required to have no less than 10 cells, otherwise the comparison was skipped. We defined genes with Benjamini-Hochberg (BH)-adjusted Wilcoxon rank sum test P-value < 0.01 and log-transformed expression fold change > 0.25 as positive markers. We took an equal number of top positive markers (ranked by AUC) from each of the two groups in comparison, and defined those as DE genes between a specific small intestine region and colon in one specific specimen. Genes defined as DE genes in comparison between any of the small intestine regions and colon, in any of the selected specimens were defined as DE genes between small intestine and colon. Similarly, to identify DE genes among different small intestine regions, we did pairwise comparison between small intestine regions in each specimen and took their union set. The only difference is positive markers in comparisons among the small intestine regions were defined as log-transformed expression fold change >0.1 and BH-adjusted Wilcoxon rank sum test P-value < 0.01. DE genes identified in any of the comparisons, which were also detected in tHIO cells, were defined as intestinal region-dependent genes.

We then calculated the average expression levels of intestinal region-dependent genes in each selected cell type, with each intestinal tissue of each specimen separated. We calculated the Spearman’s correlation coefficient between each of the small intestinal tissues and colon in each selected specimen to examine divergence between small intestine and colon. We calculated the pairwise Spearman’s correlation coefficient of different small intestinal tissues in each selected specimen to examine divergence between small intestine regions. As a background of inter-region comparison, we did pairwise Spearman’s correlation coefficient calculations between developing duodenum specimens of neighboring ages. Specifically, the day 80 old specimen was compared to the day 72 and day 85 old specimens; the day 101 old specimen was compared to the day 85 and day 122 old specimens; the day 132 old specimen, which has the most mature intestinal tissues in this study, was compared to the day 122 and day 127 old specimens. We then compared the distribution of observed small intestine - colon divergence, as well as small intestine region divergence, to the background. Significant regional difference was defined as one-sided Student’s t test nominal P < 0.01, and with more than 5 DE genes identified as described above.

We then calculated the average expression levels of intestine region-dependent genes in tHIO in each selected cell type, and compared it to its cell type counterpart of each developing intestine region of each specimen by calculating Spearman’s correlation coefficients. Median values of comparison to different specimens were presented.

### Transcriptome similarity based mapping between datasets

To map in vitro HIO, CDX2-KO and control organoid cells to the mouse gastrulation cell types, we first identified 3,000 highly variable genes in the mouse gastrulation atlas [33] using the default vst function implemented in the Seurat package. We retrieved human-mouse orthologous gene information from Ensembl (version 92). Highly variable genes with one-to-one human-mouse orthologs that were detected in the query data of human organoids were used as the feature genes. We used the average expression levels of the feature genes in each annotated mouse cell type as the reference, and calculated Spearman correlation coefficients between organoid cells and mouse cell type. Organoid cells were mapped to the mouse cell type with the highest similarity.

To map mesenchymal cells of in vitro HIO and tHIO to the developing human duodenum mesenchymal subtypes, we firstly identified developing duodenum mesenchymal subtype markers as feature genes. We used average expression levels of feature genes in each duodenum mesenchymal subtype as the reference, and calculated Spearman correlation coefficients between expression profiles of organoid mesenchymal cells and developing duodenum mesenchymal subtypes. An organoid mesenchymal cell was mapped to the duodenum mesenchymal subtype with the highest similarity.

To map non-proliferative mesenchymal cells in CDX2-KO and control organoid to the in vitro HIO non-proliferative mesenchymal subtypes, we identified markers of non-proliferative mesenchymal subtypes in 4-week in vitro HIOs as feature genes. We used average expression levels of feature genes in each of the selected subtypes as reference, and calculated Spearman correlation coefficients between CDX2-KO/control organoid non-proliferative mesenchymal cells and 4-week in vitro HIO subtypes. Selected CDX2 dataset cells were mapped to the 4-week in vitro HIO mesenchyme subtype with highest similarity. We annotated mesenchymal clusters (c1, 3, 8, 10) observed in the time course HIO data based on enrichment of 4-week in vitro HIO mesenchyme subtypes, and further mapped the CDX2 dataset cells to time course in vitro HIO mesenchymal cells accordingly.

### Identification of in vitro HIO epithelial cells that have acquired intestinal identity

To identify the in vitro HIO cells that have acquired intestinal identity, we took several filtering steps. Here, we only considered cells of day 0 (spheroid) to day 30 HIO, excluding cells of definitive endoderm. First, we calculated the Pearson’s correlation coefficients between HIO cells and cell types of mouse gastrulation (MG) stage atlas [33] using the 2,000 highly variable genes defined in the MG data, and kept the HIO clusters with minimum 80% of the cells being most similar to gut. Within these HIO clusters, HIO cells not mapping to gut were also excluded. Then we sub-clustered each sample. For samples of days 0, 3 or 7, we excluded clusters showing depleted expression of CDX2. For samples of day 14 or later, we inferred the developing organ identity based on the kNN method as described for each HIO cell. We choose day 14 as a boundary age because from day 14 on, we observed clusters enriched in cells mapping to the developing intestine. HIO samples younger than day 14 were considered as too young to use the developing human multi-organ cell atlas to determine organ identity. For samples of day 14 or later, cell clusters with more than 50% cells that were not mapping to the developing intestine, as well as any cells not mapping to developing intestine, were excluded. Finally, we examined the HIO clusters defined for the whole in vitro HIO dataset again, and further excluded the clusters with more than 75% cells excluded in the previous steps.

### Identification of differentially expressed genes during maturation of developing human duodenum epithelial stem cells

To study maturation of developing human duodenum epithelial stem cells, we ordered those cells in a pseudo-temporal order to generate a pseudotime course. We applied DiffusionMap (implemented in destiny package, with default setting except for k=20) [89] to CSS of selected cells, and used the rank of the first diffusion component as pseudotime of selected cells.

To identify genes showing differential expression levels along the constructed pseudotime course, we grouped every 50 cells into a bin based on their estimated pseudotime, and calculated the average gene expression level of every gene. Next, we applied the ‘ns’ function (implemented in the splines R package) to construct a natural spline linear regression model (df = 5) for each gene, using pseudotime orders of bins as variables and average gene expression levels of bins as response. Then we used an F-test to compare the residual of variations that could not be explained by the natural spline regression model to that of a constant model. Genes with significant expression changes along pseudotime were defined as F-test Benjamini and Hochberg (BH)-corrected P < 0.01 and log-transformed fold change of maximum versus minimum bin average expression levels > 0.5.

### Quantification of stem cell maturation and identification of stem cell maturation associated genes

We quantified stem cell maturation using a transcriptome deconvolution approach. Here we selected only epithelial cells of in vitro HIOs that have acquired intestine identity, tHIO and the developing duodenum stem cells for this analysis. To quantify the fraction of HIO spheroid and the most mature (day 132) developing duodenum epithelial stem cell identity of each selected cell, we performed hierarchical clustering to genes with significant expression changes during maturation of developing human duodenum epithelial stem cells, and selected the gene clusters showing a monotonous increase or decrease along the trajectory as feature genes for deconvolution. We then calculated the average expression level of feature genes in HIO c12 (the day 0 epithelium primed cluster) and stem cells of day 132 (the latest time point of the developing duodenum sample in the dataset), separately. Finally, we used the obtained average expression matrix as a reference transcriptome, and performed transcriptome deconvolution for each selected cell using quadratic programming via the quadprog package in R. The estimated fraction of day 132 intestinal stem cell identity, or ISC identity for short, was used as the proxy of intestinal stem cell maturity.

To identify genes related to stem cell maturation from spheroids to mature developing duodenum stem cells, we combined two different approaches. In the first approach, we firstly calculated Spearman’s correlation coefficients between ISC identity fraction and gene expression level, and then took the top 100 genes showing the strongest positive and negative correlation, respectively. In the second approach, we ordered cells included in the analysis based on their ISC identity in an ascending order. Next, the same approach as described above was applied to identify genes with expression associated with the ISC identity. We then classified genes into gene modules with hierarchical clustering on their expression patterns across bins, took the cluster that turned on at late HIO stage and kept stable expression throughout developing duodenum stage and selected 100 genes showing the highest correlation with ISC identity fraction in this cluster.

For identification of differentially expressed genes between the developing and adult duodenum stem cells, we selected the developing stem cells with 100% ISC identity fraction, and compared to adult duodenum stem cells using the presto R package. Genes with significant differential expression levels between developing and adult stem cells were defined as AUC > 0.6, log-transformed expression fold change > 0.25 and expressed in over 25% cells of the group being tested.

### Inference and visualization of between cell type interaction

We applied CellPhoneDB (v2.0) [80] to normalized expression levels of epithelial and mesenchymal cell types of the developing duodenum samples with default parameters. Only epithelial cell types with more than 50 cells, and mesenchymal cell types with more than 100 cells were examined. We defined significant interactions as BH-corrected P < 0.05. To visualize cell type interactions, we connected each cell type to 5 other cell types with the highest number of interactions, and visualized the interaction network with the igraph R package in the undirected mode using the Fruchterman-Reingold layout.

### Pseudo-spatial reconstruction of scRNA-seq data

To reconstruct the pseudo-spatial distribution of tHIO epithelial cells as well as the developing duodenum epithelial and mesenchymal cells, we applied novoSpaRc [51] to the corresponding scRNA-seq data sets. For the developing duodenum, we used the day 132 tissue sample. We first obtained the top 100 cell type markers (ranked by AUC, with presto R package) for each cell type in epithelium and mesenchyme, separately. We used the normalized expression levels of the identified markers as well as 22 reference genes as the inputs for novoSpaRc. The chosen reference genes are APOA4, BMP3, CDH1, CHGA, COL1A2, DLL1, EGF, EPCAM, ERBB3, F3, GPX3, LGR5, MKI67, MUC2, NPY, NRG1, PDGFRA, RSPO2, RSPO3, SLC2A2, TAGLN and WNT2B. We manually curated binarized expression patterns of selected reference genes, according to previously reported in situ hybridization (RNAscope) experiments in the developing human intestine [14, 30] and mouse enterocyte zonation atlas [52]. Binarized expression patterns of reference genes were used to guide novoSpaRc inference. Reference intestine morphology with epithelium and mesenchyme was provided to novoSpaRc. novoSpaRc was run with the author suggested parameters (https://github.com/rajewsky-lab/novosparc).

For tHIO, we used 4-week and 8-week tHIO epithelial cell types that committed to intestinal fate. The chosen reference genes were MKI67, CDH1, EPCAM, SLC2A2, APOA4, LGR5, MUC2 and CHGA. Reference intestine morphology with epithelium was provided to novoSpaRc. Other procedures are the same as those for the developing duodenum sample.

### Construction of transcription factor (TF) co-expression network

We used HIO clusters 0, 1, 2, 3, 5, 7, 8, 9, 10, 12, 14 to construct the TF co-expression network associated with the development of intestinal epithelium and mesenchyme. Non-intestine or non-mesenchyme HIO clusters were removed in this analysis. The Human TF list was downloaded from Human TFDB [90]. With differential expression analysis done by the presto R package, we identified cluster enriched TFs with the criteria AUC > 0.6, log-transformed fold change >0.1 and expressed in more than 10 cells. We calculated Pearson’s correlation coefficients of cluster enriched TF expression patterns across clusters. Only correlation > 0.8 was defined as a significant correlation, and only genes with no less than 10 expression-correlated genes were kept. We then used the igraph for network visualization.

### Curation of GI disease-associated genes

The list of disease genes was drawn from the NIH Genetics Home Reference (https://ghr.nlm.nih.gov) and The Genetic and Rare Diseases Information Center (GARD) (https://rarediseases.info.nih.gov). We included all genes they listed as being associated with a GI disease from these two databases. When there are no genes reported to be associated with a GI disease listed in these resources, we manually searched for at least one human study reported on the particular GI-disease. The curated disease-associated genes are listed in Table S7. Median expression levels of NPSR1 and NLRC4 in each tHIO cell type were 0, so these two genes are not presented in the heatmap of Figure S12. All other genes were listed in the Table S7.

## Supplemental Figures

**Figure S1.**
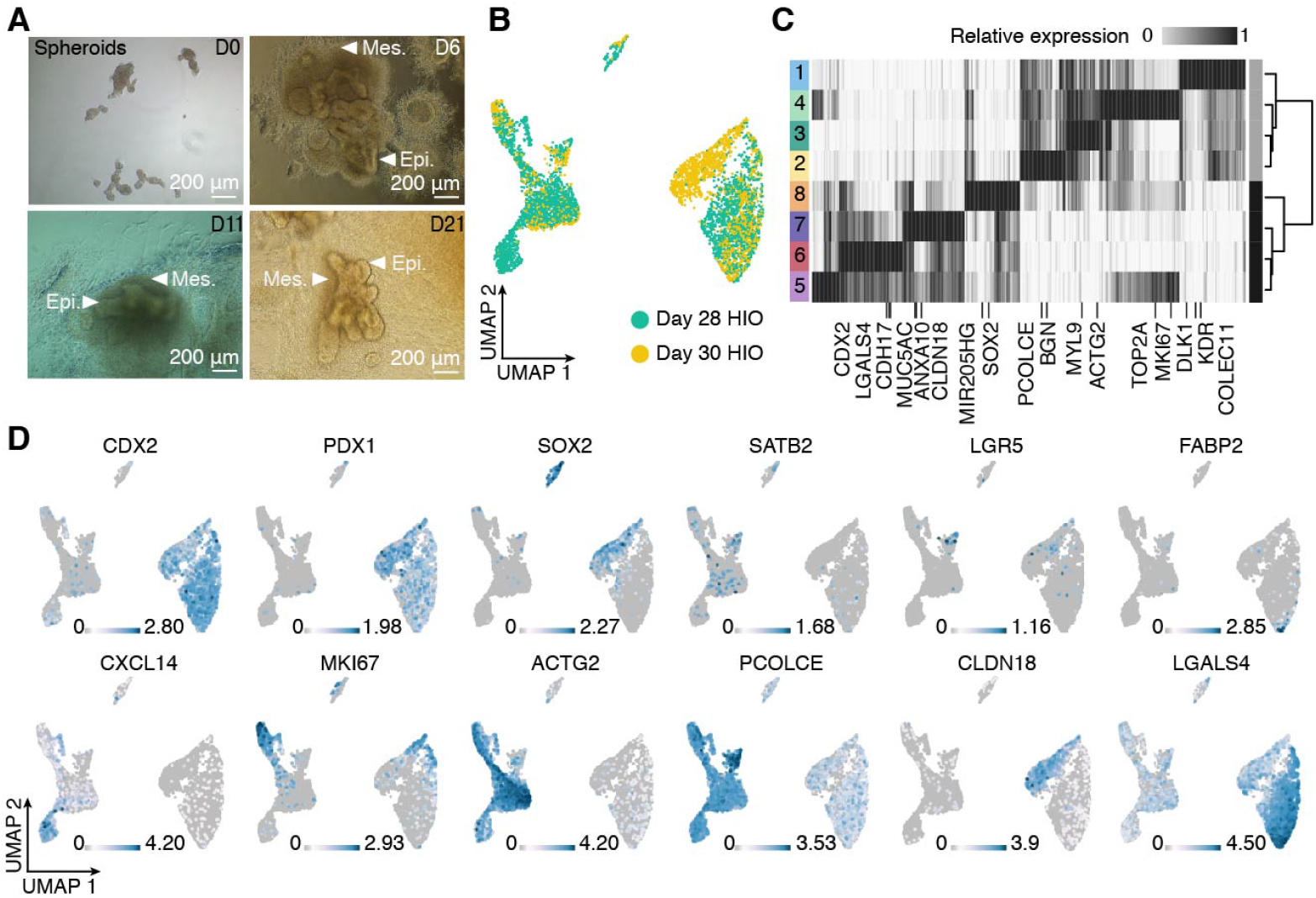
Histological and cell composition analysis of in vitro HIOs. A) Brightfield images at day 0 (spheroid), 6, 11, 21 of HIO differentiation. Mesenchyme can be observed developing around bulging epithelium. Scale bar, 200 μm. B) UMAP embedding of d28 and d30 HIO scRNA-seq data, with cells colored by time point/batch. C) Heatmap shows cluster average expression of marker genes (column) for each HIO cluster (row). Sidebar denotes epithelial (dark grey) and mesenchymal (light grey) clusters. D) Feature plots show representative marker gene expression.

**Figure S2.**
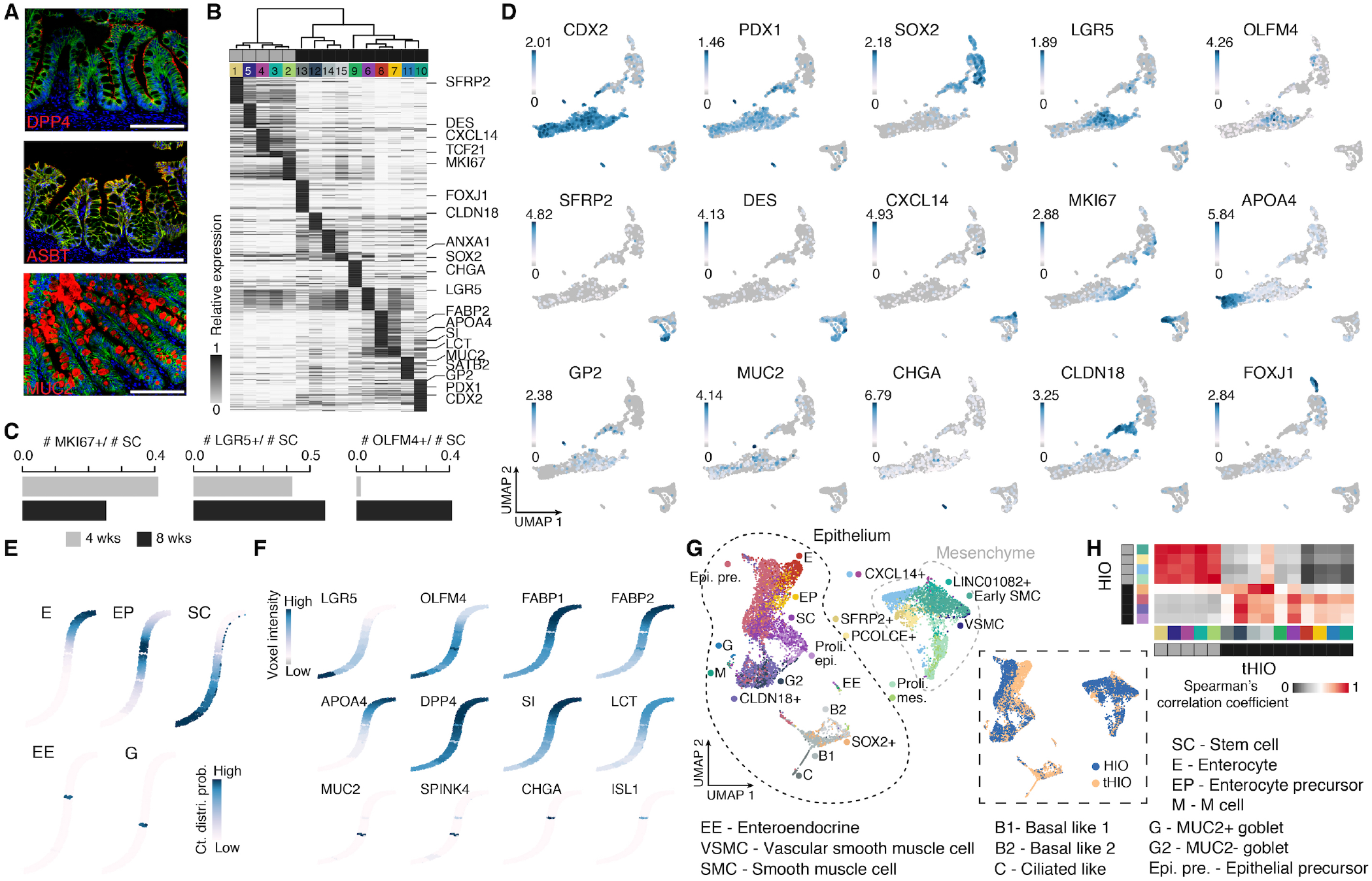
Cell composition analysis of transplanted HIOs, epithelial pseudo-spatial reconstruction, and integration with in vitro HIOs. A) tHIO histological analysis approximately 10 weeks after transplantation into an immunocompromised mouse kidney capsule. Dipeptidyl Peptidase 4, DPP4; Mucin 2, MUC2; Apical Sodium–Bile Acid Transporter, ASBT. Scale bars, 200 μm. B) Heatmap shows cluster average expression of marker genes (row) for each tHIO cluster (column) with selected genes shown. C) Bar plots show the proportion of cells within the stem cell cluster (c6) that express MKI67, LGR5, or OLFM4 in 4- or 8-week tHIOs. D) Feature plots showing representative marker gene expression across tHIO cells. E-F) Feature plots showing the cumulative cell distribution probability of each cell type (E) or marker gene expression (F) in a pseudo-spatial reconstruction (novoSpaRc) of the epithelium based on the reference in situ hybridization data from developing duodenum [30, 52]. Enterocyte, E; Enteroendocrine, EE; MUC2+ goblet, G; Enterocyte precursor, EP; Intestinal stem cell, SC. G) Harmony integrated UMAP embedding of HIOs (28- and 30-day) and tHIOs (4- and 8-week) with cells colored by cell type (left) or source (right). H) Heatmap shows the transcriptome similarity between HIO and tHIO cell types. Sidebar shows cell groups colored by mesenchyme (light grey) or epithelium (dark grey), and by cluster.

**Figure S3.**
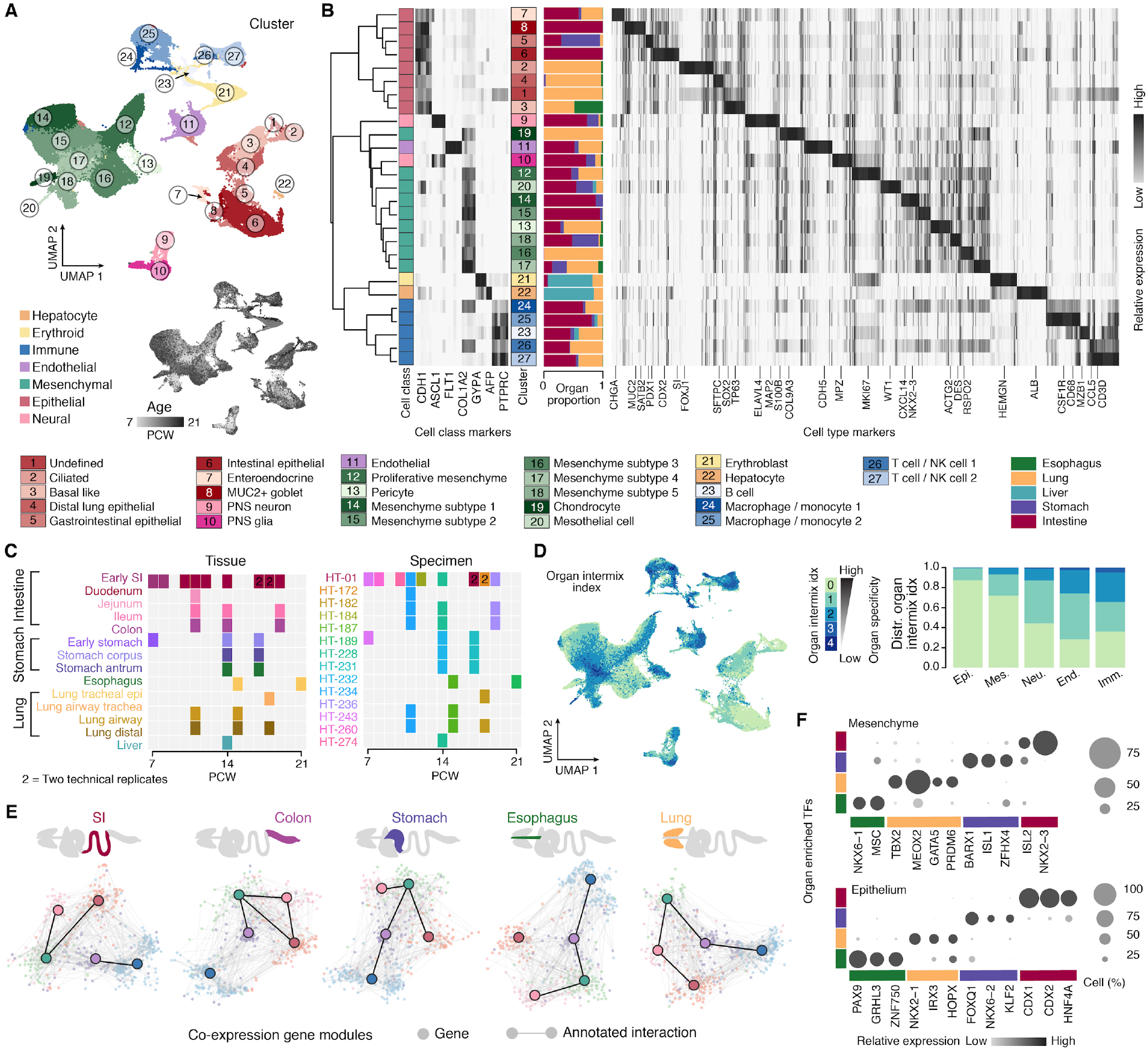
Characterization of the developing human multi-organ cell atlas. A) UMAP embedding of the developing human multi-organ atlas scRNA-seq data with clusters indicated and colored by major cell class. Inset shows cells colored by age. B) The dendrogram shows cluster similarity (Pearson) based on highly variable genes, with selected class markers, top cluster marker genes, and a sidebar showing the proportion of cells of each organ per cluster. C) Tissue (left) and individual specimen (right) information of the developing tissues used in this work. D) Left, UMAP embedding of the developing human reference atlas colored by organ intermixing index. This index represents the number of organs with shared transcriptome features of each cell, excluding the organ of the examined cell. See also STAR methods. Right, stacked bar plots show the distribution of organ intermixing index of each cell class. E) Interacting gene network of each organ with layout based on receptor and ligand co-expression. Each receptor or ligand encoding gene was colored by cell class with the maximum expression level. Grey lines represent annotated receptor-ligand pairs [80]. Dark lines between cell classes indicate more annotated interacting gene pairs than expected by chance. F) Dot plots show the expression of organ enriched transcription factors (TF) in the mesenchyme (top) or epithelium (bottom).

**Figure S4.**
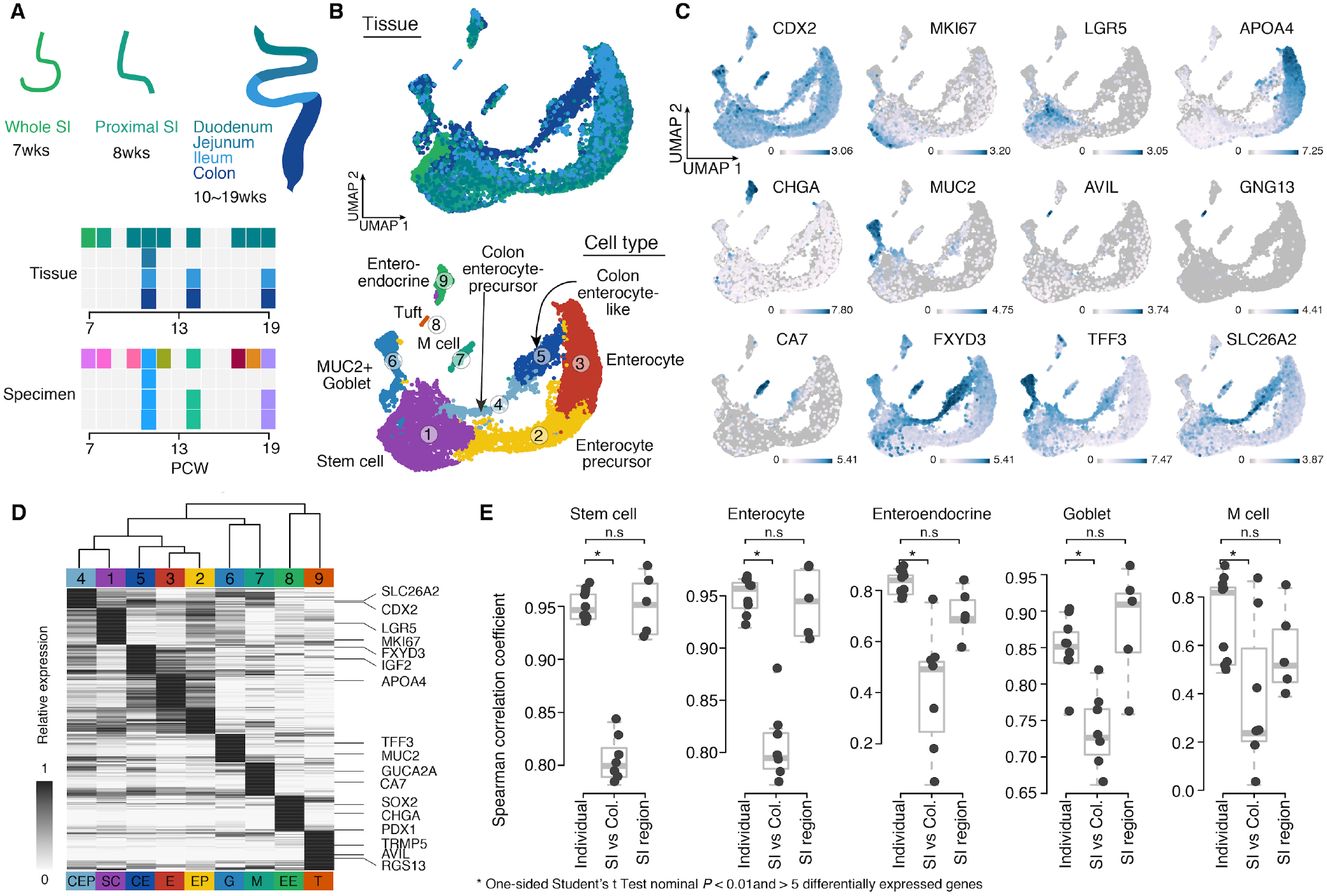
Inter-regional analysis of the developing human intestine epithelium. A) Region and time point information for each developing intestine tissue sample analyzed in this study. B) CSS-integrated UMAP embedding of the developing intestine epithelial cells colored by region (top) or cell type (bottom). C) Feature plots of intestine epithelial cell type markers. D) Heatmap shows relative average expression levels of cell type markers, with selected genes highlighted. E) Every three box plots show transcriptome similarity between duodenum samples of neighboring ages (Individual), between different small intestine regions and colon (SI vs Col.), or between different small intestine regions of the same cell type (SI region). A comparison between different tissues was only performed using tissues from the same individuals. Genes showing differential expression levels between intestine regions were identified in each epithelial cell type separately, and used to calculate Spearman’s correlation coefficients.

**Figure S5.**
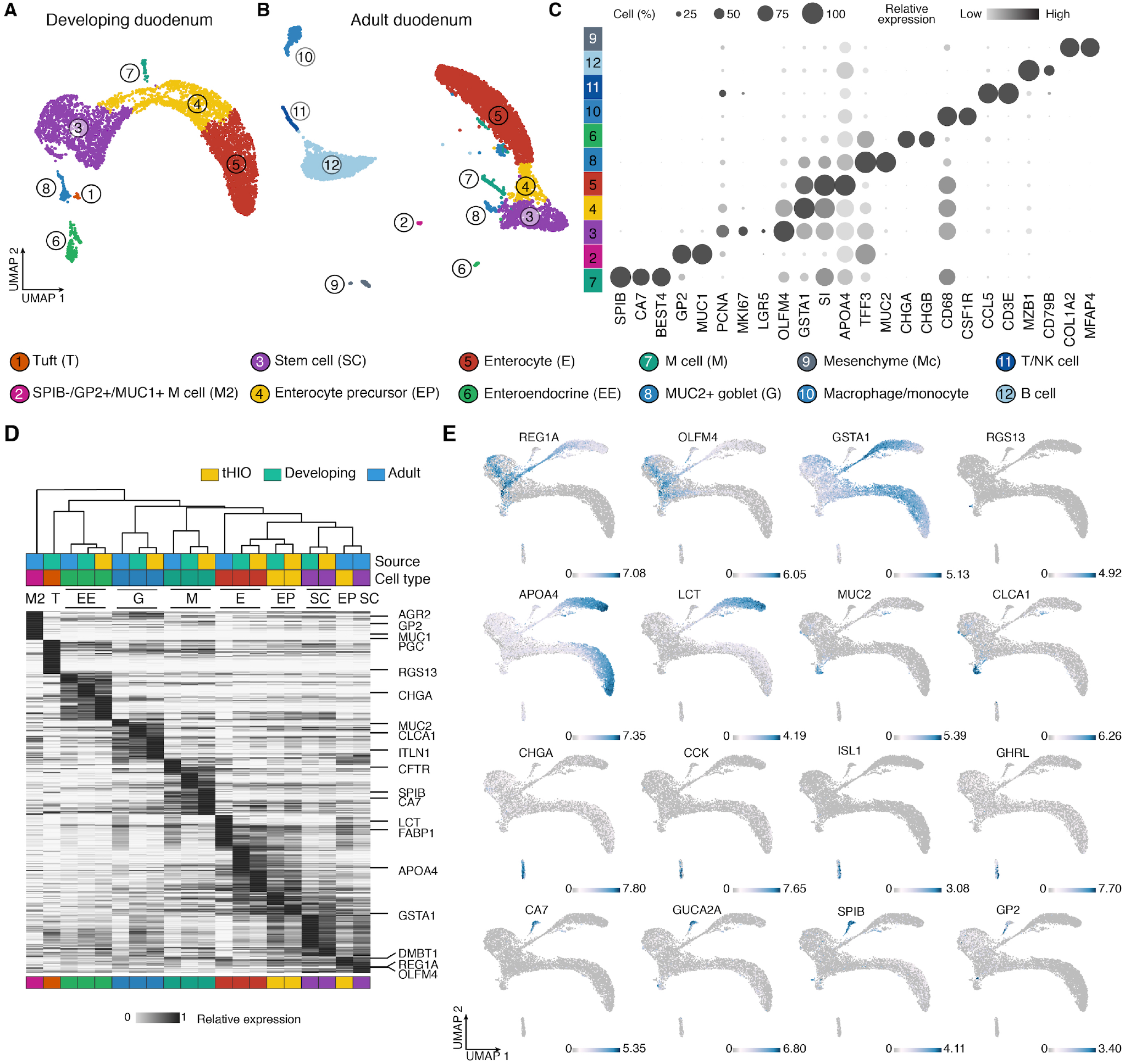
Integrative analysis of tHIO and human primary small intestine epithelium. A-B) UMAP embedding of the developing duodenum (A) and adult (B) epithelial cells colored and numbered by annotated cell type. C) Dot plot shows the average gene expression levels (color) and expressed proportion (size) of cell type markers in adult duodenum. D) Heatmap shows relative cluster average expression of intestinal epithelial cell type marker genes in tHIO, developing and adult intestine integrated data. E) Feature plots show cell type marker expressions in the tHIO, developing and adult intestine integrated UMAP embedding.

**Figure S6.**
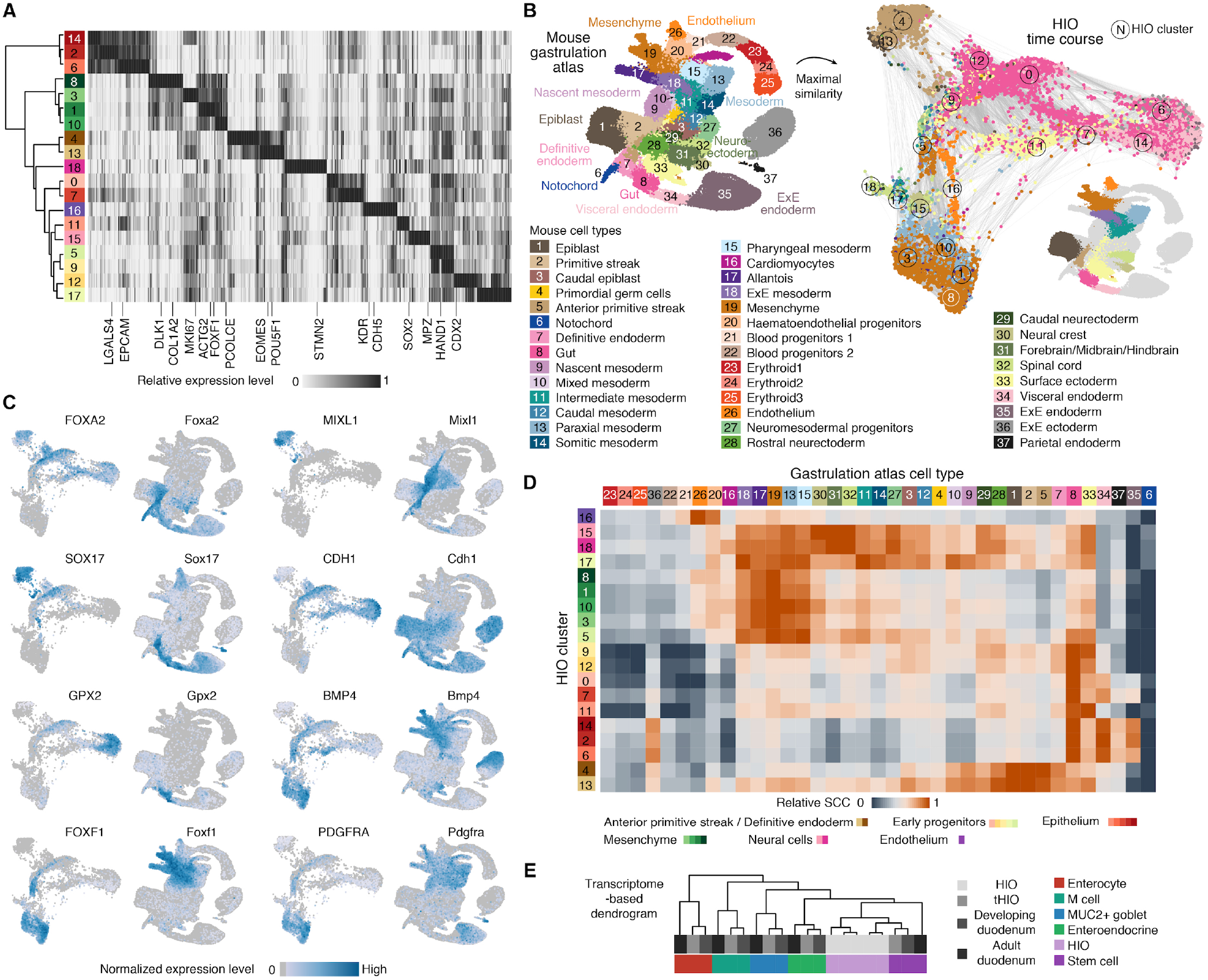
Mouse gastrulation map enables assessment of cell composition of in vitro HIO time course data. A) Heatmap showing relative expression levels of top 50 cluster markers across each HIO time course cluster. B) HIO cells were compared to a single-cell reference atlas covering multiple stages of mouse gastrulation (MG) [33]. HIO SPRING plot colored by maximum Spearman correlation coefficient (SCC) to the MG reference cluster. Inset shows MG clusters with similarity in the HIO. C) Feature plots of definitive endoderm (FOXA2 and SOX17), primitive streak (MIXL1), gastrointestinal epithelium (CDH1 and GPX2) and mesoderm or mesenchyme (BMP4, FOXF1 and PDGFRA) markers in HIO time course and MG data. D) Heatmap shows the transcriptome similarity (Spearman’s correlation coefficients, SCCs) between HIO and MG clusters using highly variable genes in the MG dataset that have expressed human one-to-one orthologs. E) Hierarchical clustering of average transcriptome Pearson’s correlations between the developing, adult, and tHIO cell type counterparts together with HIO epithelial clusters committed to intestine identity, using cell type markers identified in either developing or adult duodenum.

**Figure S7.**
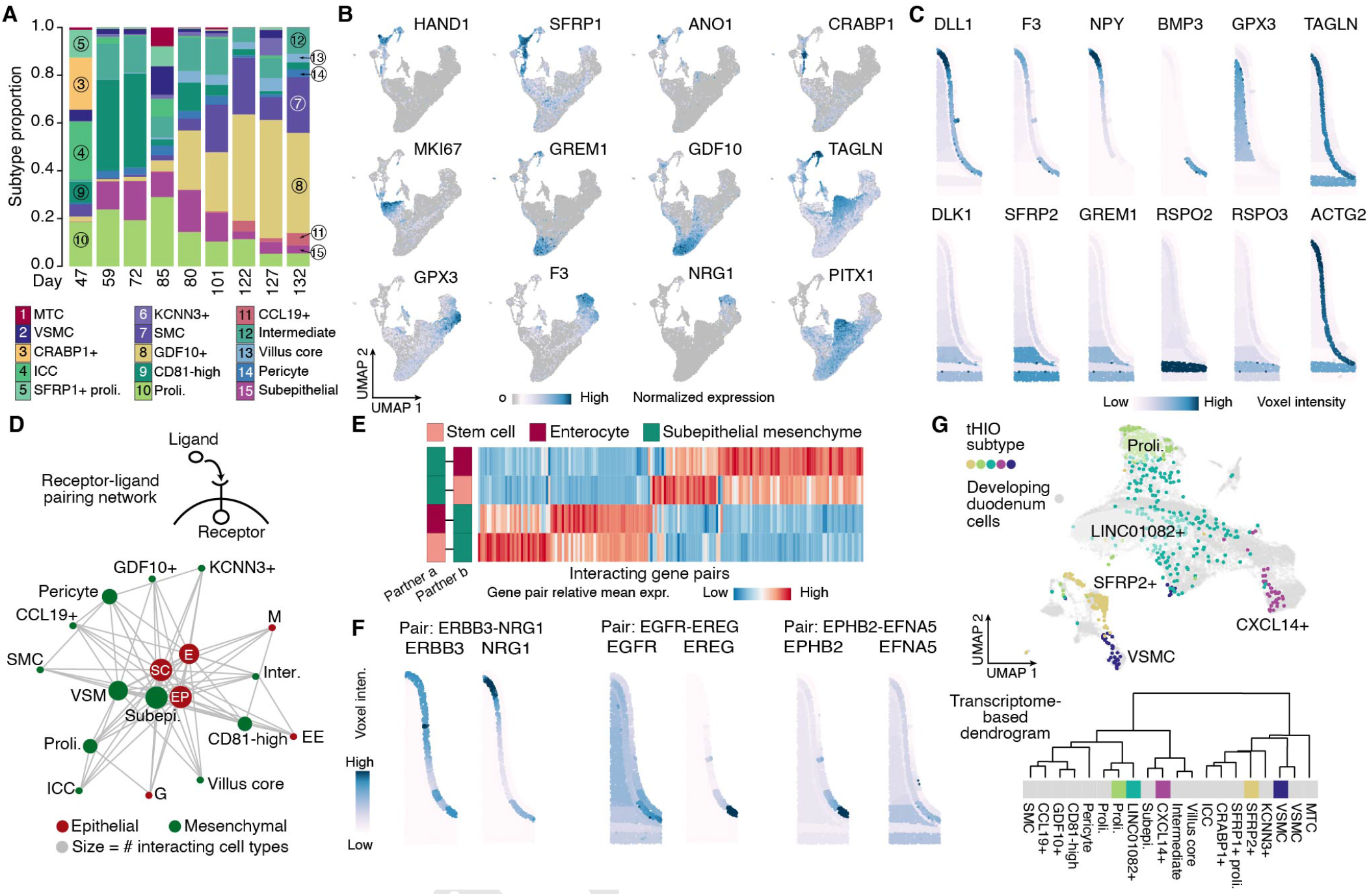
Temporal progression of developing human duodenal mesenchyme and predicted interactions with epithelium. A) The stacked bar plot shows the proportion of mesenchymal subtypes at different time points. B) Feature plots of duodenal UMAP embedding showing expression of mesenchymal subtype marker genes. C) Feature plots showing mesenchymal subtype marker gene expression on the pseudo-spatial epithelium and mesenchyme space reconstructed with novoSpaRc and in situ hybridization data [30, 52]. D) Epithelial and mesenchymal cell type interaction network inferred by CellPhoneDB according to annotated interacting receptor and ligand gene pair expression levels. Each cell subtype is linked with five other most frequently linked subtypes. E) Heatmap shows relative mean expression levels of significant receptor-ligand pairing interactions between stem cell or enterocyte and subepithelial mesenchyme. F) Feature plots of gene pairs that show a significant interaction between stem cell and subepithelial mesenchyme on the reconstructed pseudo-spatial space. G) Top, Integrated UMAP embedding of mesenchymal cells from the developing duodenum and tHIO. tHIO cells are colored by de novo identified cluster identity shown in Figure 1; grey cells represent developing human duodenum cells. Bottom, hierarchical clustering of tHIO and the developing duodenum mesenchymal subtypes based on cluster average expression of subtype markers.

**Figure S8.**
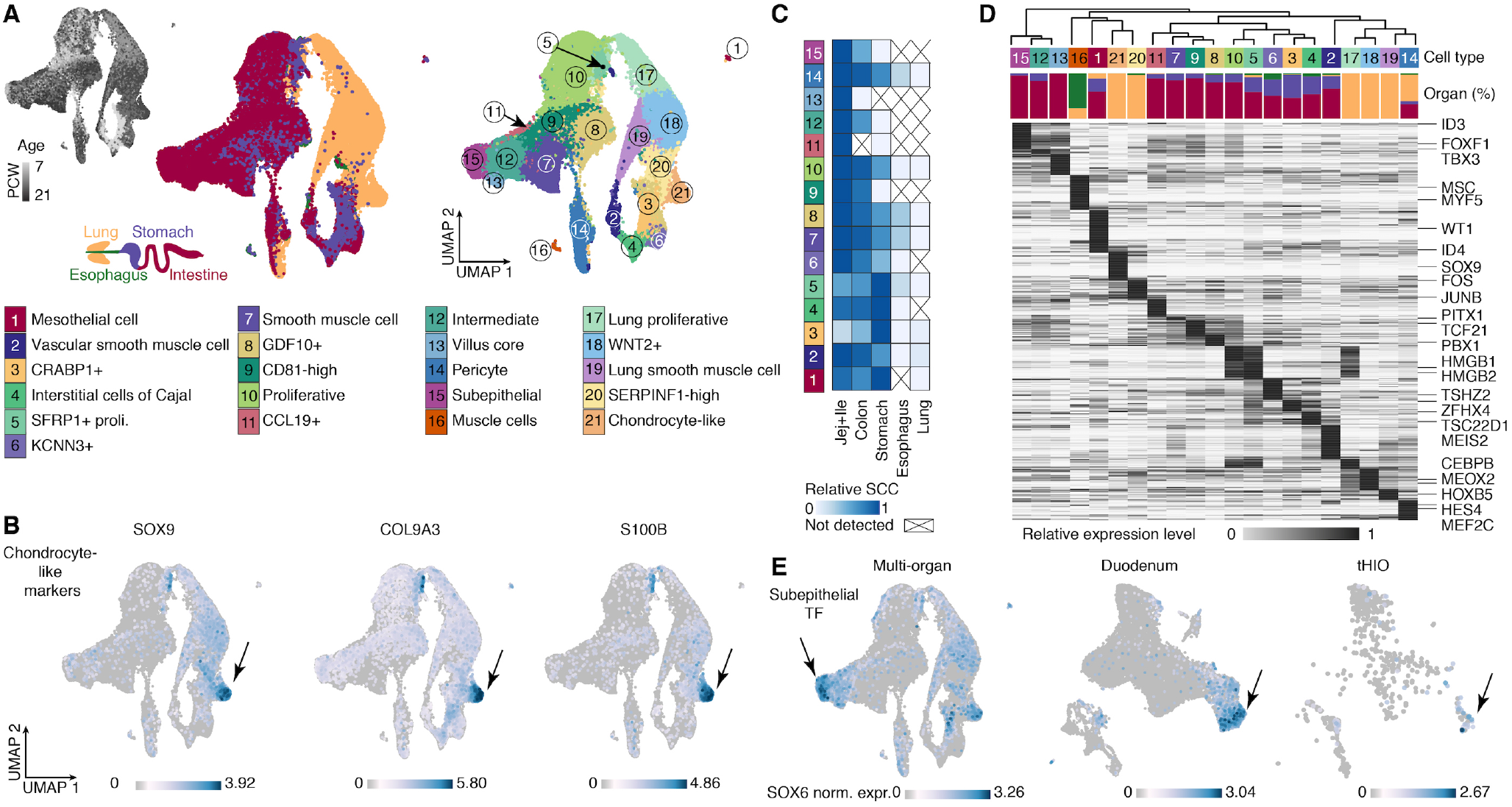
Identification of features that are specific to mesenchymal subtypes in the developing multi-organ atlas. A) UMAP embedding of mesenchymal cells in the developing human reference atlas colored by (from left to right) sample age, organ, and cell subtype. B) Feature plots of the chondrocyte-like mesenchymal subtype (c21) markers. C) Heatmap shows the Spearman’s correlation coefficient quantifying the transcriptome similarity of subtype counterparts between the duodenum and other small intestine regions or organs. D) Heatmap shows relative cluster average expression levels of subtype markers. Transcription factors are highlighted. The transcriptome-based dendrogram was constructed by correlation distance of highly variable genes. E) Feature plots of SOX6 in mesenchymal cells of the developing human multi-organ reference, developing duodenum reference, and tHIO data. Cell embedding of the developing duodenum and tHIO mesenchyme are the same as shown in Figure 5F.

**Figure S9.**
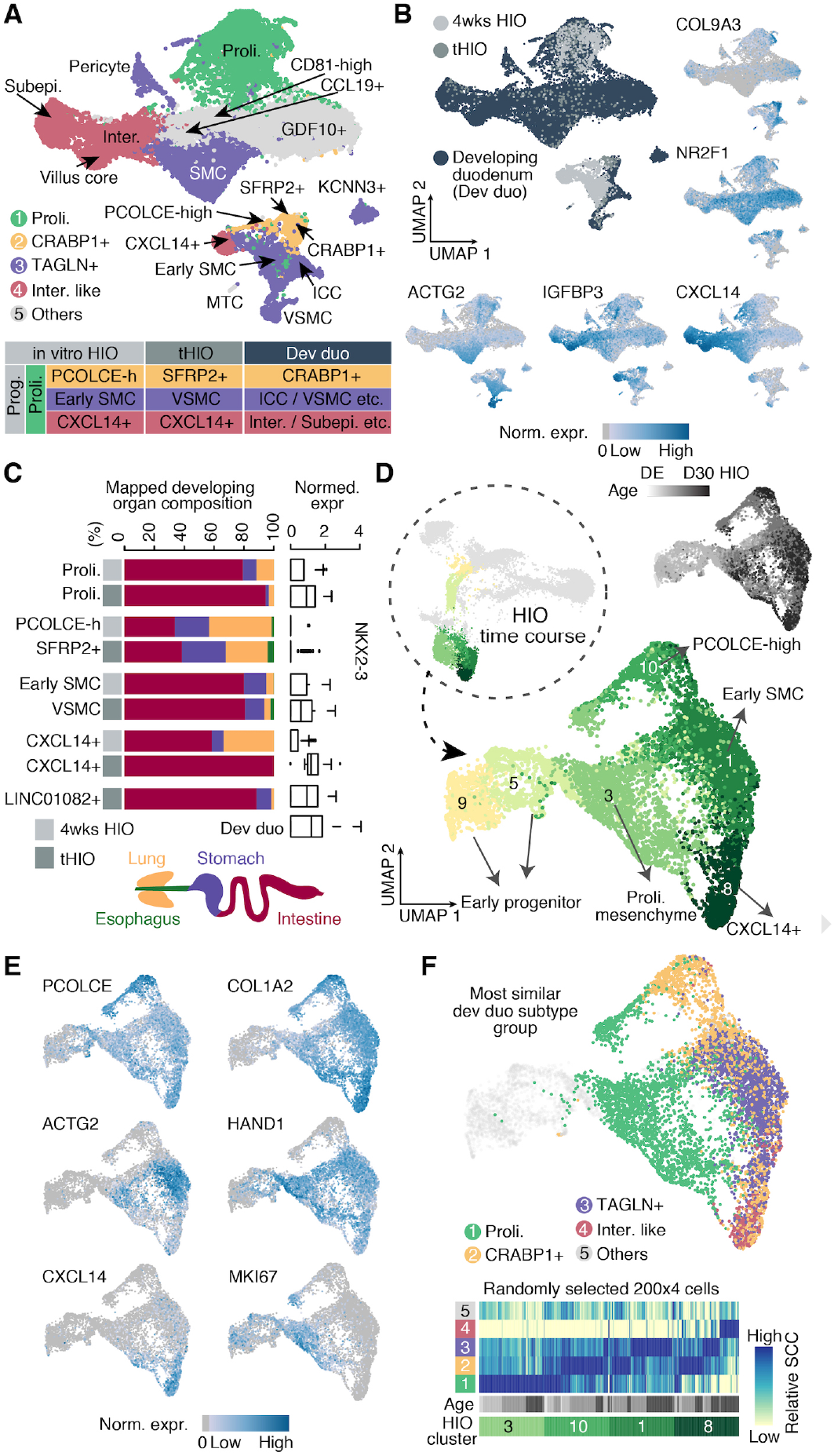
Reconstruction of mesenchymal subtype development over the HIO in vitro time course. A) Integrated UMAP embedding of 4 week-HIO, tHIO and the developing duodenum mesenchymal cells, with cells colored by developing duodenum subtype groups and inferred counterparts in the in vitro HIOs and tHIOs. B) Integrated UMAP with cells colored by source, or expression of differentially expressed genes that support the inferred subtype correspondence between different sources. COL9A3+/NR2F1 + mark PCOLCE-high (PCOLCE-h, in vitro HIOs), SFPR2+ (tHIOs) and CRABP1+ (developing duodenum); IGFBP3+/CXCL14+ mark CXCL14+ (in vitro HIOs), CXCL14+ (tHIOs) and subepithelial/intermediate/villus core (developing duodenum); ACTG2 mark early smooth muscle cell (early SMC, in vitro HIOs), vascular smooth muscle cells (VSMC, tHIO) and ACTG2+/TAGLN+ populations (developing duodenum). C) Stacked barplot shows the proportion of cells in subtypes of 4-week HIO (light grey) and tHIO (darker grey) mapping to organs of the developing human atlas. The box plot shows NKX2-3 expression in each organoid subtype and median of all developing duodenum subtypes. D) In vitro HIO time course mesodermal/mesenchymal cells were extracted (top left, clusters from original SPRING embedding are highlighted) and projected to a new UMAP embedding to visualize mesenchymal cell developmental trajectories. Cells are colored, numbered, and annotated by HIO time course cluster (bottom), or age (top right). E) Feature plots show expression distributions of differentially expressed genes. F) In vitro HIO mesenchymal cells were compared to the developing duodenum subtypes. Top, UMAP embedding with HIO mesenchymal cells colored by the most similar developing duodenum mesenchymal subtype groups. Transcriptome similarities were quantified as Spearman’s correlation coefficient (SCC), which was calculated with top cluster markers of each developing duodenum subtype. Subtype groups were defined as subtypes with similar transcriptome. Bottom, the heatmap shows similarity patterns between the randomly selected in vitro HIO cell subset and each developing duodenum subtype group. For groups combining multiple subtypes, the maximum similarity is presented.

**Figure S10.**
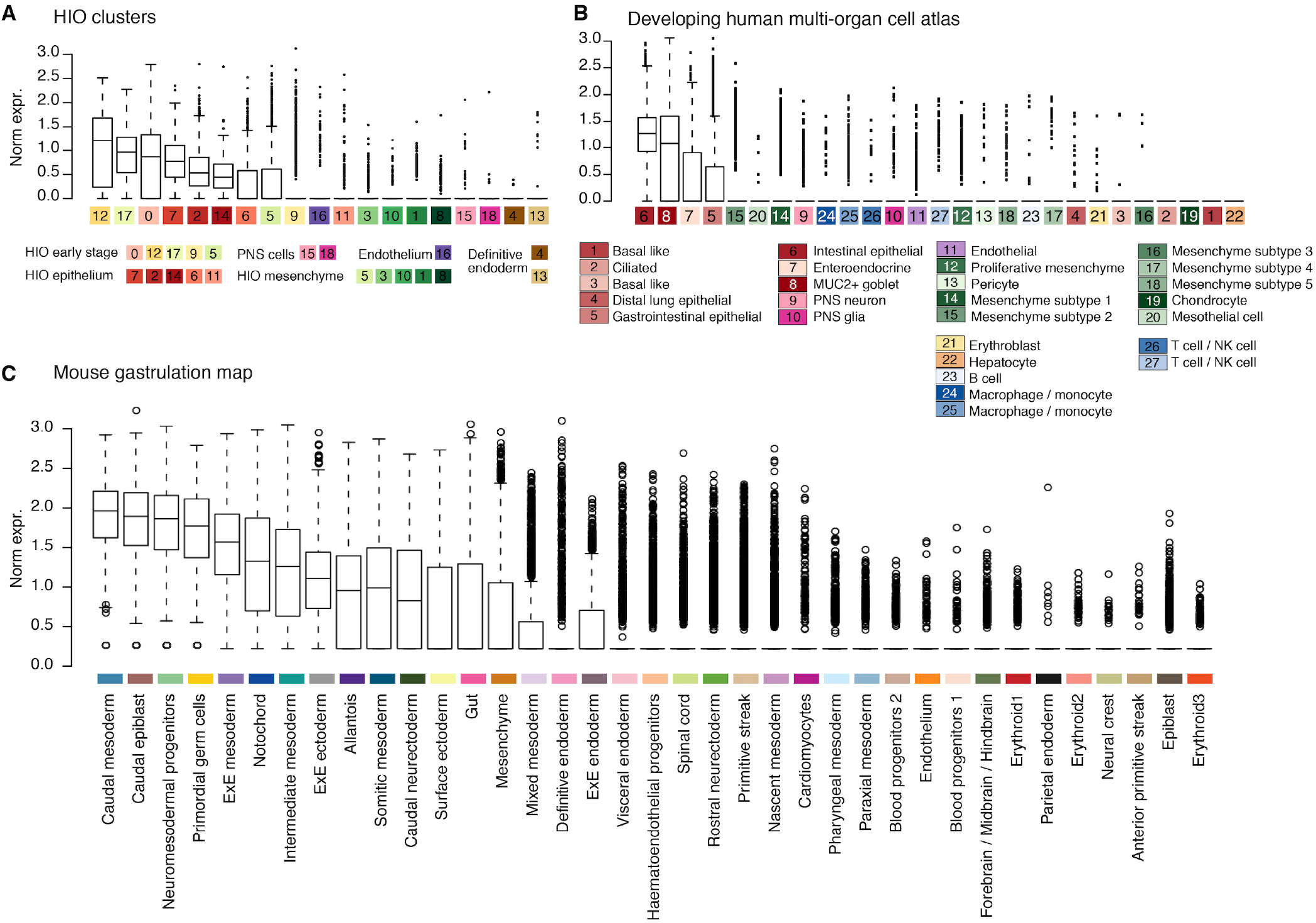
CDX2 expression across reference maps. A-C) Boxplots show expression of CDX2 across HIO time course (A), developing human multi-organ cell atlas (B), and mouse gastrulation atlas (C) cell clusters.

**Figure S11.**
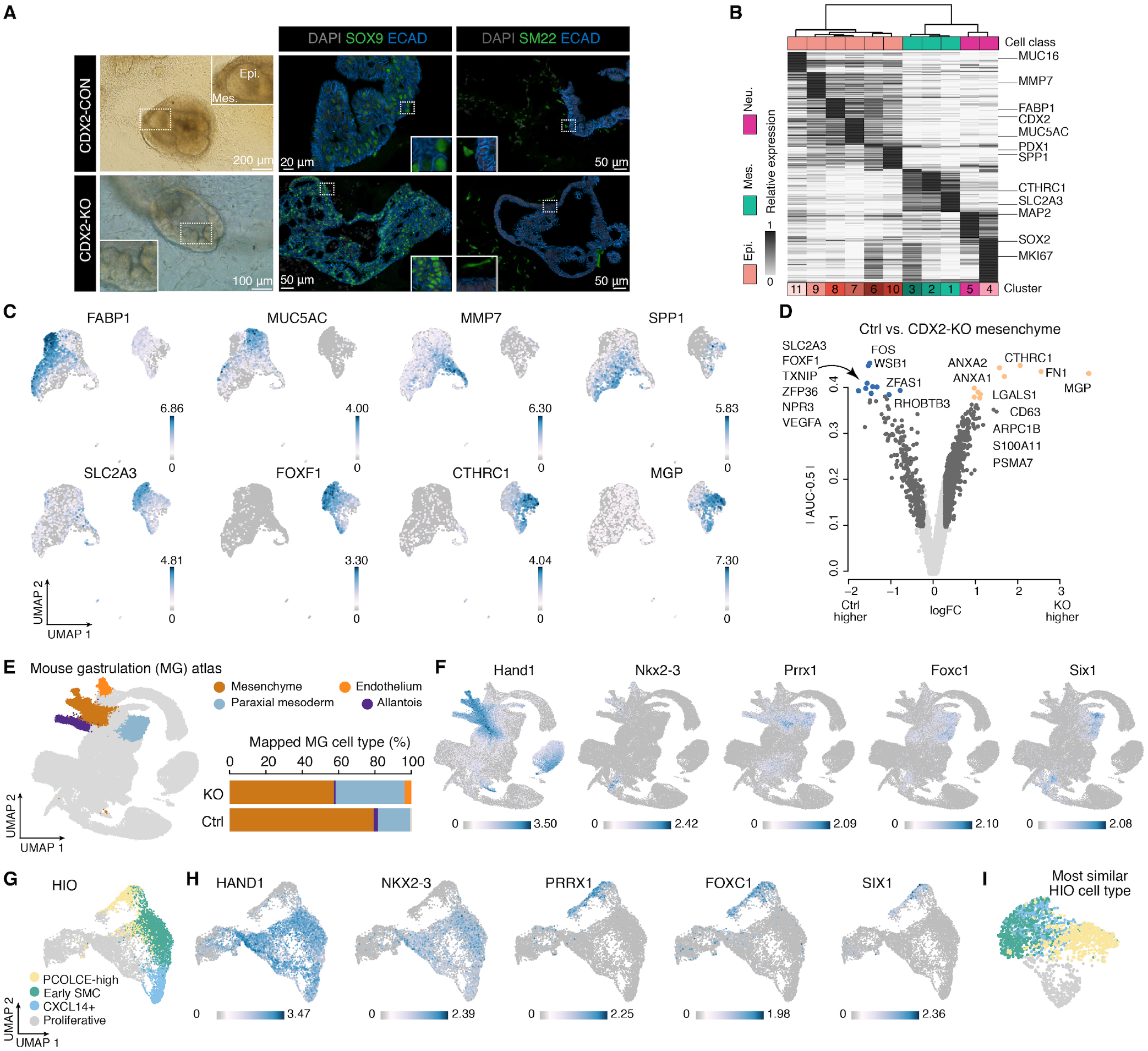
Cell composition and differential gene expression analysis of control and CDX2-knockout (KO) PSC-derived organoids. A) Bright-field and immunofluorescence imaging of control and CDX2-KO PSC-derived organoids. B) Heatmap shows relative cluster average expression levels of marker genes (row) of each cluster (column) of the CDX2 dataset. C) Feature plots of cell cluster marker genes on the integrated UMAP embedding shown in Figure 7. D) Volcano plot shows differentially expressed genes between CTHRC1+ cells of CDX2-KO organoids and SLC2A3+ cells of control organoids. Dark grey indicates genes with significant differences between groups. Yellow and blue indicate the top 10 positive markers of each group. E) Left, UMAP of published mouse gastrulation (MG) atlas, with mesenchymal cell types mapped to CDX2 dataset highlighted. Right, stacked bar plot shows the proportion of mesenchymal cells mapped to MG cell types. F) MG atlas feature plots of TFs showing differential expression levels between control and CDX2-KO organoid mesenchyme. G) UMAP embedding of in vitro HIO time course mesenchymal progenitor/mesenchymal cells (c1/3/5/8/9/10 from Figure 4B) with cells colored by cell type assignment. Same UMAP embedding as shown in Figure S9D. H) Feature plots of control and CDX2-KO mesenchyme differentially expressed transcription factors (TFs) in the HIO time course mesoderm/mesenchyme cells. I) UMAP embedding of CDX2 dataset mesenchymal cells colored by most similar HIO mesenchymal subtypes.

**Figure S12.**
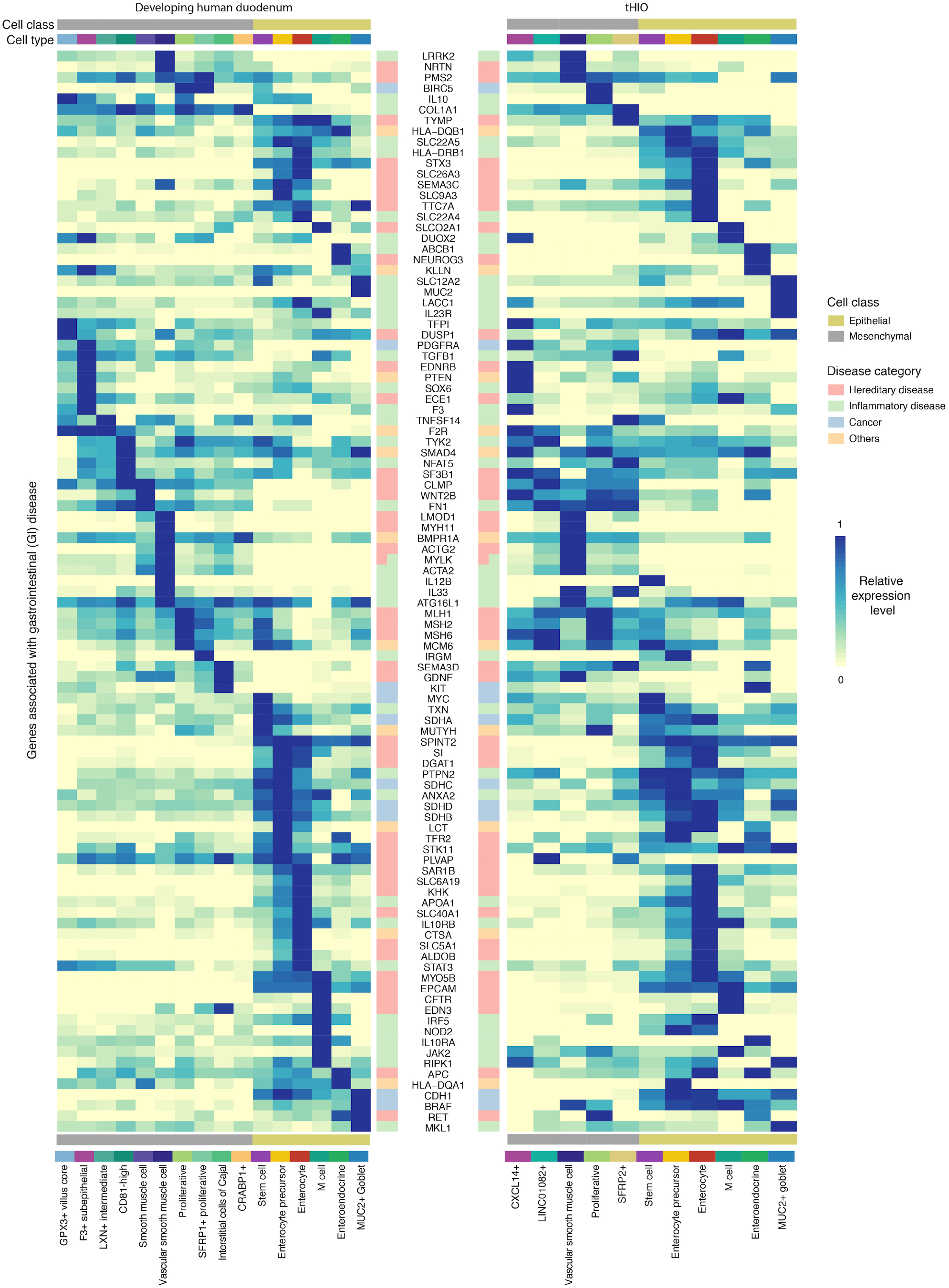
Expression of gastrointestinal (GI) disease-associated genes. Heatmap shows average expression levels of GI disease-associated genes in the developing duodenum (left) and tHIO (right) cell types. Each row represents a gene and each column represents a cell type. The procedure of manual curation of GI disease-associated genes is in the STAR methods section.

## Supplemental Tables

**Table S1: Sample information and scRNA-seq data quality check.**

The table presents information of samples used in this study, as well as cell number, median detected gene number, medium UMI number and medium percentage of transcripts mapped to the mitochondrial of each sample after filtering.

**Table S2: Cell type markers.**

Table present top cell type markers of 4-week HIO, 4- and 8-week tHIO, adult duodenum, CDX2-dataset, developing human multi-organ cell atlas, the developing duodenum mesenchyme, the developing duodenum epithelium, the developing intestine, HIO time course and multi-organ mesenchyme.

**Table S3: List and GO enrichment of genes differentially expressed between the developing duodenum epithelial stem cell and adult counterparts.**

**Table S4: List and GO enrichment of genes associated with intestine epithelial stem cell emergence and maturation.**

**Table S5: Interactions between mesenchyme and epithelium in the developing duodenum.**

The table presents nominal P values and mean expression of interacting pairs between the developing duodenum epithelial and mesenchymal subtypes, as well as interacting gene pairs with Benjamini Hochberg corrected P < 0.05 between subepithelial mesenchyme and stem cell.

**Table S6: CDX2-knockout induced differentially expressed genes.**

The table presents CDX2-knockout induced top 50 differentially expressed genes in epithelium and mesenchyme separately.

**Table S7: Manually curated gastrointestinal disease-associated genes.**

## Supplemental Videos

3D UMAP embedding of epithelial cells of tHIO and duodenum, with cells colored by cell types, source and stem cell scores, respectively.

